# Unsupervised approach to decomposing neural tuning variability

**DOI:** 10.1101/2022.03.19.484958

**Authors:** Rong Zhu, Xue-Xin Wei

## Abstract

Neural representation is often described by the tuning curves of individual neurons with respect to certain stimulus variables. Despite this tradition, it has become increasingly clear that neural tuning can vary substantially in accordance with a collection of internal and external factors. A challenge we are facing is the lack of appropriate methods to accurately capture trial-to-trial tuning variability directly from the noisy neural responses. Here we introduce an unsupervised statistical approach, Poisson functional principal component analysis (Pf-PCA), which identifies different sources of systematic tuning fluctuations, moreover encompassing several current models (e.g.,multiplicative gain models) as special cases. Applying this method to neural data recorded from macaque primary visual cortex– a paradigmatic case for which the tuning curve approach has been scientific essential– we discovered a novel simple relationship governing the variability of orientation tuning, which unifies different types of gain changes proposed previously. By decomposing the neural tuning variability into interpretable components, our method enables discovery of new structure of the neural code, capturing the influence of the stimulus drive and internal states simultaneously.

## 1 Introduction

A central goal in neuroscience is to determine how the neural responses depend on external stimulus variables and the internal states of the brain. The dependence of individual neuron’s firing rate on certain stimulus variable is often described by the turning curve, i.e., the average firing rate of neuron as a function of the stimulus [1, 2, 3, 4]. Because tuning curves are the consequences of various internal computations in neural circuits, it is likely and indeed empirically the case that it could be modulated by factors other than the stimulus variable selected *a priori* [5, 6, 7, 8, 9, 10, 11, 12, 13, 14, 15, 16, 17, 18, 19, 20, 21]. Indeed, variability of neural tuning has now been widely reported in neural systems, and furthermore been proposed to exhibit various forms, including multiplicative gain [22, 6, 23, 24, 25, 18, 26, 27], additive gain change [5, 28, 29, 30, 31], shift of tuning peaks [32, 33, 34], and tuning width changes [35]. These observations reflect the role of various factors, whether related to stimuli (e.g. stimulus contrast [36], stimulus history [37, 33]), behavior (e.g. movement [16]), or latent brain states [25, 28].

Tuning variability has been widely implicated both functionally, i.e., information encoding and the behavioral performance [38, 15, 28, 30, 39, 19], and mechanistically, i.e., how tuning variability is generated [40, 41, 42]. Prior studies have attempted to quantitatively model the variability of tuning in sensory cortex, in particular orientation tuning in primary visual cortex (V1), which is widely considered as a paradigmatic case for studying neural code. Decades of studies in V1 have shed general insights regarding how neurons in cortex encode external sensory variables. Perhaps surprisingly, studies of V1 tuning variability have yielded results that are seemingly at odds so far. One line of work [25, 26] has proposed a simple multiplicative gain model to account for the tuning variability. Multiplicative gain has been postulated to have a vital role for encoding contrast [43], encoding uncertainty [27], facilitating downstream readout [44], implementing attention [6, 7], as well as the transformation of coordinate systems (e.g. retina- to body-centered) in parietal cortex [45, 46]. Mechanistic models suggest that multiplicative gain could result from threshold-linear neurons operating in the presence of intrinsic intracellular noise [40, 41, 42]. In contrast to the above work, other studies have suggested additive interactions [5, 47, 18] or both additive and multiplicative gain [28, 30, 31] in V1.

Crucially, the analysis methods in the majority of this prior work presumed relatively restrictive structure for tuning variability (*e*.*g*.,[25, 18, 26, 28, 30]), leaving open the question of whether other forms of fluctuations might in fact account for the data better. Furthermore, existing analyses generally relied on trial-averaging and comparison across conditions [6, 37, 30], thus failing to capture the trial-to-trial variability in tuning. Addressing these open issues requires approaches that can infer the structure of tuning fluctuations directly on single-trial data - and ideally on the raw spike train itself - while also avoiding restrictive assumptions.

Here we introduce a new unsupervised statistical technique, Poisson functional PCA (Pf-PCA), to identify the structure of of latent tuning fluctuations directly from neural spiking data. Importantly, we apply this method to address tuning variability in a classic neural system that has long been characterized via tuning, namely neurons in V1. Because Pf-PCA yields a generative model of the trial-to-trial tuning variability, it could be used to analyze the information encoding through the calculation of information-theoretical measures such as Fisher information. It could also be used to analyze the geometrical structure of the neural manifold. Performing these analyses, we find that Pf-PCA reveals several new insights. The proposed analysis framework is broadly applicable to other low-dimensional tuning modalities.

## 2 Results

### 2.1 The Poisson functional PCA framework

Fig. 1a illustrates the basic modeling framework of Pf-PCA (see Method for details). The model assumes that the logarithm of the tuning curves (of an arbitrary stimulus variable) is determined by a *smooth mean component* and *smooth* functional principal components (fPCs) weighted by the amount of latent fluctuations. Note that each fPC is a function that is tuned to the stimulus variable. The fPCs and their weights (i.e., scores) together capture the fluctuations of the tuning curves for individual trials. Quantitatively, the tuning curve ***µ***_*t*_ for the *t*-th trial can be describe as

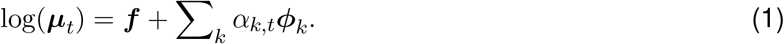

**Figure 1:**
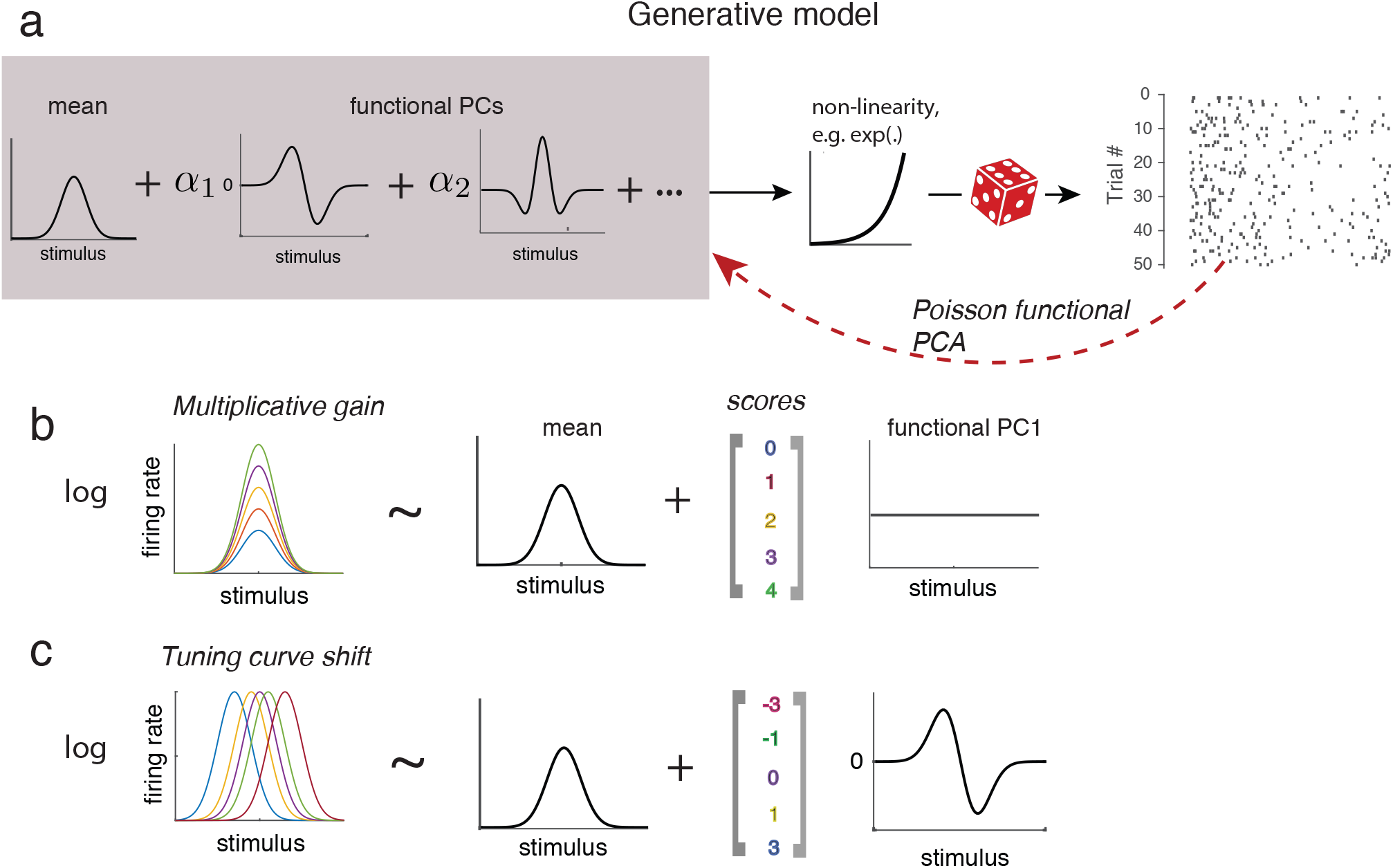
The Poisson functional PCA (Pf-PCA) framework. (a) Illustration of the Pf-PCA framework. Tuning functions (i.e., the logarithm of the tuning curves) are modeled as a sum of the mean and functional principal components (fPCs) weighted by the amount of latent fluctuations *α* (i.e., scores). Tuning functions then pass through a static non-linearity to obtain the tuning curves. The non-linearity is assumed to be an exponential function in this study. Finally, standard Poisson spiking noise is assumed for the generation of the observed spike trains. The Pf-PCA algorithm performs inference on the observed spike counts. It extracts the mean component, the fPCs, and the trial-to-trial scores of each fPC. (b,c) Schematics showing that this framework could account for multiplicative gain and tuning shift as two special cases.

Here ***f*** is the *mean component*, ***ϕ***_*k*_ is the *k*-th fPC, *α*_*k,t*_ denotes the amount of fluctuation (i.e., score) for the *k*-th component during the *t*-th trial, and is assumed to follow a zero-mean Gaussian distribution. The spike train on every trial is assumed to be generated from a Poisson process with the firing rate specified by Eqn. (1). Note that with only the first term ***f***, this model is equivalent to the standard tuning curve model of spike counts. The second term Σ_*k*_ *α*_*k,t*_***ϕ***_*k*_ is the additional variance from the contribution of trial-to-trial fluctuations, making the model naturally capture the over-dispersion of spike counts.

Our algorithm takes the spike count data as the input and infers the mean, fPCs, the variance of each components, as well as the weight for each fPC for each trial. Critically, the shape of fPCs, which specifies the particular form of the fluctuations, is directly inferred from the data. When studying the neural response to continuous stimulus variables, it is natural to assume the mean component and fPCs of individual neurons as some smooth functions of stimulus. Importantly, our method merely assumes that the mean component and fPCs are smooth, without imposing restrictive assumptions on their shapes. Our method provides a novel way to parse the variability of tuning into a set of fPCs from the spike counts. Now consider a couple of special cases. With the additional assumptions that there is only one fPC and that it is constant over the stimulus dimension (Fig. 1b), our model becomes essentially the multiplicative gain model [25]. When tuning curves exhibit systematic lateral shift, our model could capture it with a fPC that is proportional to the derivative of the mean component (Fig. 1c). It is worth emphasizing that our method is general, and capable to capture other cases that may be potentially more complicated. It may be used to analyze the tuning fluctuations of any one-dimensional stimulus variables with smooth tuning properties.

Our method is developed by adapting a technique, *i*.*e*., functional PCA [48, 49, 50, 51, 52], to deal with Poisson spiking noise. As the firing rate is not observed, it makes the inference procedure more challenging. We resolve this problem by developing a procedure based on an Expectation-Maximization algorithm. Details of the inference procedure are described in the Method section. In broad strokes, the algorithm treats the unobservable firing rates *µ*_*t*_(*s*) as “observations” generated from the model in Eqn. (1). To maximize the likelihood, the algorithm iterates between estimating the mean and covariance matrix parameters and calculating their posteriors based on the spike counts given their current estimates via Monte Carlo method. This step gives an estimate of the firing rates. In the next step, we apply the functional PCA technique to the estimated firing rates for estimating the components and the trial-to-trial fluctuations.

### 2.2 Validation of the method

We validated our method systematically using simulated data. Inspired by previous experimental observations on tuning variability [22, 34, 33, 23, 32, 35, 25, 28, 30], we first examined whether our method is able to recover fPCs that correspond to additive gain change, multiplicative gain change, tuning shift, or sharpening. Specifically, we generated synthetic data, which exhibit different types of tuning fluctuations by reverse-engineering the appropriate fPC, and tested Pf-PCA and alternative methods with these data, where the ground-truth were known.

Fig. 2a shows results based on the analysis of the simulated datasets using our method and alternative methods (see Method for details). We found that Pf-PCA could accurately recover the form of the fluctuations in all four cases. Furthermore, it approximately recovers the proportions of variance explained by the structured fluctuation (Fig. 2a), as well as the magnitude of the latent fluctuation on a trial-by-trial basis.

**Figure 2:**
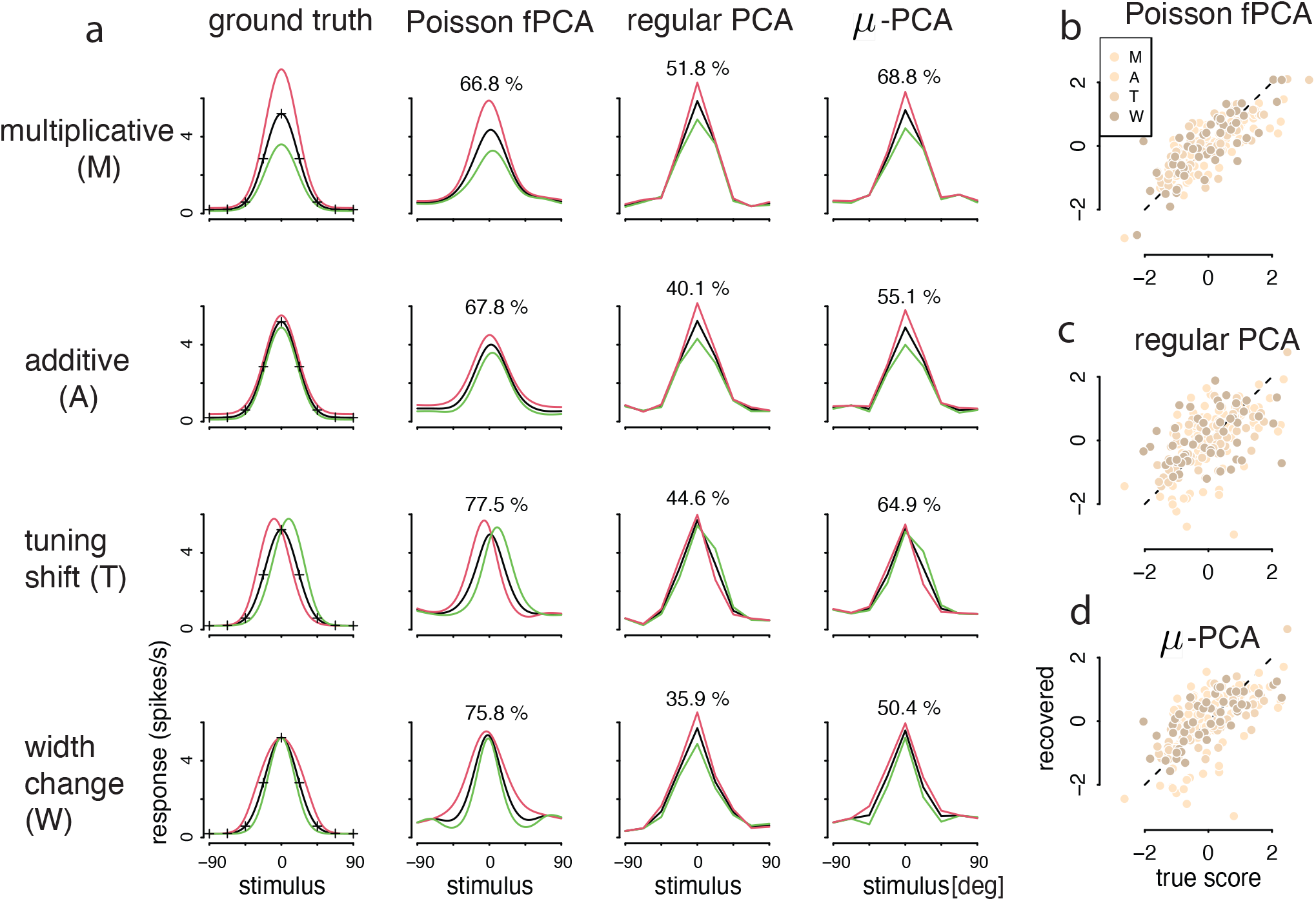
Model validation with bell-shape tuning curves. (a) Inferred fluctuations of four types of tuning fluctuations. From left to right: ground truth, Pf-PCA, regular PCA, and a reduced version of Pf-PCA termed as *µ*-PCA. In each case, we simulated neural responses from the corresponding ground truth model, and then inferred the form of the fluctuation and its magnitude for each trial. The percentages on the plots show the proportion of variance explained by the component. By our construction of the model, the first fPC of a “perfect” estimation procedure should explain 80% of the variance. (b,c,d) Recovered scores v.s true scores for each method. Light to dark bisque points denote multiplicative gain (“M”), additive gain (“A”), tuning shift (“T”), and tuning sharpening (“W”). Dashed lines indicates the diagonal. Overall, Pf-PCA substantially outperforms the alternatives in all these cases, both in terms of recovering the form of the tuning fluctuations and the magnitude on each trial. The correlations between the recovered and the ground truth are 0.788 for Pf-PCA (b), 0.544 for regular PCA (c), and 0.647 for *µ*-PCA (d).

How does our method compare to simpler methods? Applying conventional PCA to the synthetic data, we found that it often misidentified the form of the fluctuation, and that it could not reliably estimate the magnitude of the latent fluctuations (Fig. 2b). We also applied a variant of our method, referred to as *µ*-PCA, by removing the smoothness constraint in our full algorithm (see Method for details). This algorithm is similar to the Poisson PCA [53] (a discussion of the technical differences between *µ*-PCA and Poisson-PCA can be found in SI.) The *µ*-PCA generally performs better than regular PCA, but is still considerably worse than the full method Pf-PCA.

We further validated our method when multiple types of fluctuations co-exist, *e*.*g*., a combination of multiplicative gain and tuning shift. We found that Pf-PCA could recover both components reliably (see Fig. S1), and that it drastically outperforms regular PCA and *µ*-PCA (Fig. S2). Additionally, we validated our method in the case of monotonic tuning curves, *e*.*g*., the sigmoidal tuning curves, and found similar results (see Fig. S3). Taken together, these results on synthetic data suggest that our method could robustly recover the structure and magnitude of the tuning fluctuations using experimentally realistic amount of trials.

### 2.3 Pf-PCA reveals power-law modulation of neural tuning

We next show that our method could be used to reveal scientific insights into the neural codes. We will focus on the variability of orientation tuning in macaque V1, which has been a question of substantial interest in the past decades and may have general implications regarding the principle of neural coding in cortex. Previous studies have mainly focused on “gain variability”, which assumes that constant additive gain or multiplicative gain which scales the whole thing curve. The nature of this tuning variability has been heavily debated to date. Our unsupervised approach enables us to generalize the notion of “gain variability” to general “tuning variability”, resulting in a more accurate understanding of the structure of the neural response.

We analyzed seven previously published datasets, each with dozens of neurons simultaneous recorded from macaque V1 [54, 30] (total number of neurons = 402). During these experiments [54, 30], drift gratings with different directions were presented, each for 1 or 1.28 seconds. A block-randomized design was used in these experiments, with each block sampled a pre-determined set of stimulus directions once. See Method for details. To build some intuitions on orientation tuning variability, we first splitted the trials (i.e.,blocks) into two halves according to the number of spikes for individual neuron, and calculated the tuning curves for the high and low conditions [30]. Fig. 3a shows six representative example neurons. Visual inspections suggest that tuning variability is heterogeneous across neurons, exhibiting features consistent with additive gain or multiplicative gain or both, though other times neither.

**Figure 3:**
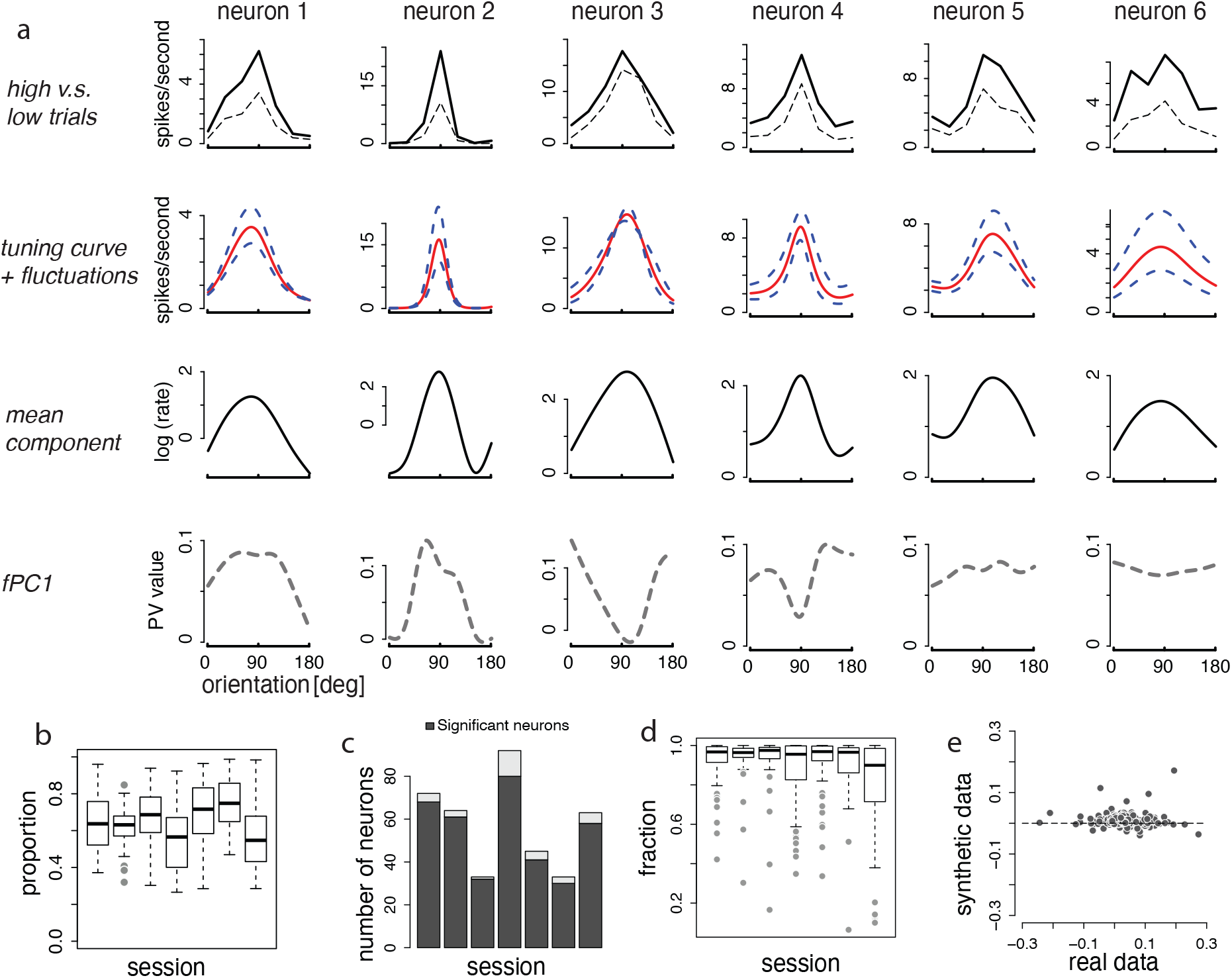
Results of Pf-PCA on V1 data. (a) Recovered mean component and first fPC for six example neurons. Neuronal tuning fluctuations exhibit a variety of structures. The top panels show the average tuning curves by splitting the trials into high and low trials. Panels in the second row show the tuning curves and the fluctuations inferred from Pf-PCA, plotted in the firing rate space. The last two rows show the mean component and the first fPC, both plotted in the logarithm of the firing rate scale. For the last row, “PV” (y-axis) represents the value of the first fPC for each stimulus. (b) The variance explained by the first fPC is typically above 60%. (c) Testing the significance of the regression analysis of the first fPC on the mean component. The majority of the neurons show a significant linear relationship between the mean component and the first fPC. (d) The percentages of the first fPC explained by the linear on the mean component in each session. We defined the “fraction” index which captures the percentage of the first fPC explained by the linear on the mean. The fact that “fraction” is closed 1 suggests that the linear function of the mean highly explains the first fPC for the majority of the V1 neurons. (e) Slope values of the regression analysis obtained from real data v.s. those obtained from synthetic data. The synthetic data were generated from a multiplicative gain modulation model.

We applied Pf-PCA to analyze the tuning fluctuations for stimulus orientation. We treated each block of stimuli as one trial, assuming that the tuning curve is stable within each block. When applying Pf-PCA, we assumed three fPCs, which are sufficient to capture most of the tuning variability in these data (Fig. S4). In fact, the first fPC alone captures 62.4% of the variance on average (Fig. 3b). Below we will focus our analysis primarily based on the first fPC.

As mentioned before, if a neuron exhibits multiplicative gain change, the first fPC should be a constant (Fig. 1b). However, we found that the first fPC for the majority of neurons is not constant (for examples, see Fig. 3a). This implies that the fluctuations of the firing rate of these neurons could not be accurately described as a pure multiplicative gain, and instead the gain appears to be stimulus-dependent. Interestingly, the first fPC for most neurons is highly correlated with the mean component. This is confirmed by a simple linear regression analysis between the mean component and the first fPC (Fig. 3c). For quantification, we defined a fraction to capture the percentage of the first fPC explained by the simple linear relationship (see Method for details), and found that this linear relationship explains the most of the information in the first fPC (Fig. 3d). We wondered if our estimation procedure might exhibit systematic biases so that even when the ground truth model was a simple gain modulation model, the estimated first fPC might nonetheless be correlated with the mean component. We performed a control analysis and found that this is unlikely. Specifically, we simulated datasets from a multiplicative gain model with approximately matched statistics and performed Pf-PCA on the synthetic datasets (see Method for details). The results showed that the slope values of the regression for the synthetic data are much closer 0 compared to what we obtained from the V1 data (Fig.3e).

Crucially, the above observation (linear relationship between the mean tuning curve and the first fPC) has conceptually important implications for tuning structure. In particular, the linear relationship permits the following linear approximation for the first fPC *ϕ*_1_(*s*),

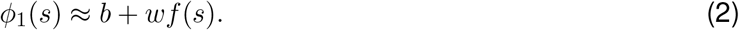

Together with Eqn. 1 and some algebraic manipulations, we found that the tuning curve for trial *t* can be expressed as

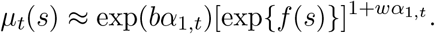

Because the latent variable *α*_1,*t*_ appears in the exponent, it suggests that the fluctuations of the tuning curves could be in fact described as a power-law modulation, with the exponent of the power function varying from trial to trial.

### 2.4 Power-law modulation accounts for both additive and multiplicative gain change

Previous studies have proposed two forms of gain change in V1 [25, 28, 29, 30], i.e. additive and multiplicative. It has been heavily debated which type of variability is more approximate to describe the V1 activity, or whether both types of activity co-exist in V1. We hypothesize that the part of controversy is due to the restrictive notion of gain variability in previous studies. By considering and analyzing general tuning variability as enabled by Pf-PCA, below we will demonstrate that the power-law relation unifies these different forms of gain variability.

Noticing *µ*_0_(*s*) = exp{*f* (*s*)} (and assuming that it is already normalized to have peak activity equal to 1 by absorbing into the intercept term *b*), the tuning curve on each trial can be re-expressed as

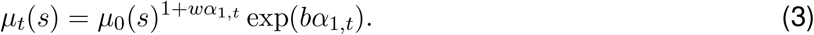

Eqn. (3) is a power function with the power 1 + *wα*_1,*t*_ and the scale *bα*_1,*t*_, both of which are linear function of the fluctuation *α*_1,*t*_ for trial *t*. In this relation, for each neuron there are two free parameters corresponding to the *slope* and *intercept* in the regression analysis respectively. Without loss of generality, we can constrain the intercept to be always non-negative. The consequence of varying each parameter on the tuning is straight-forward to see. Specifically, a non-zero intercept would lead to fluctuation of the peak firing rate, while non-zero slope would lead to systematic tuning width change due to the exponentiation (see Fig. 4a). Depending on the specific combination of the slope (*w*) and intercept (*b*), the tuning fluctuation will exhibit different characteristics for individual neurons.

**Figure 4:**
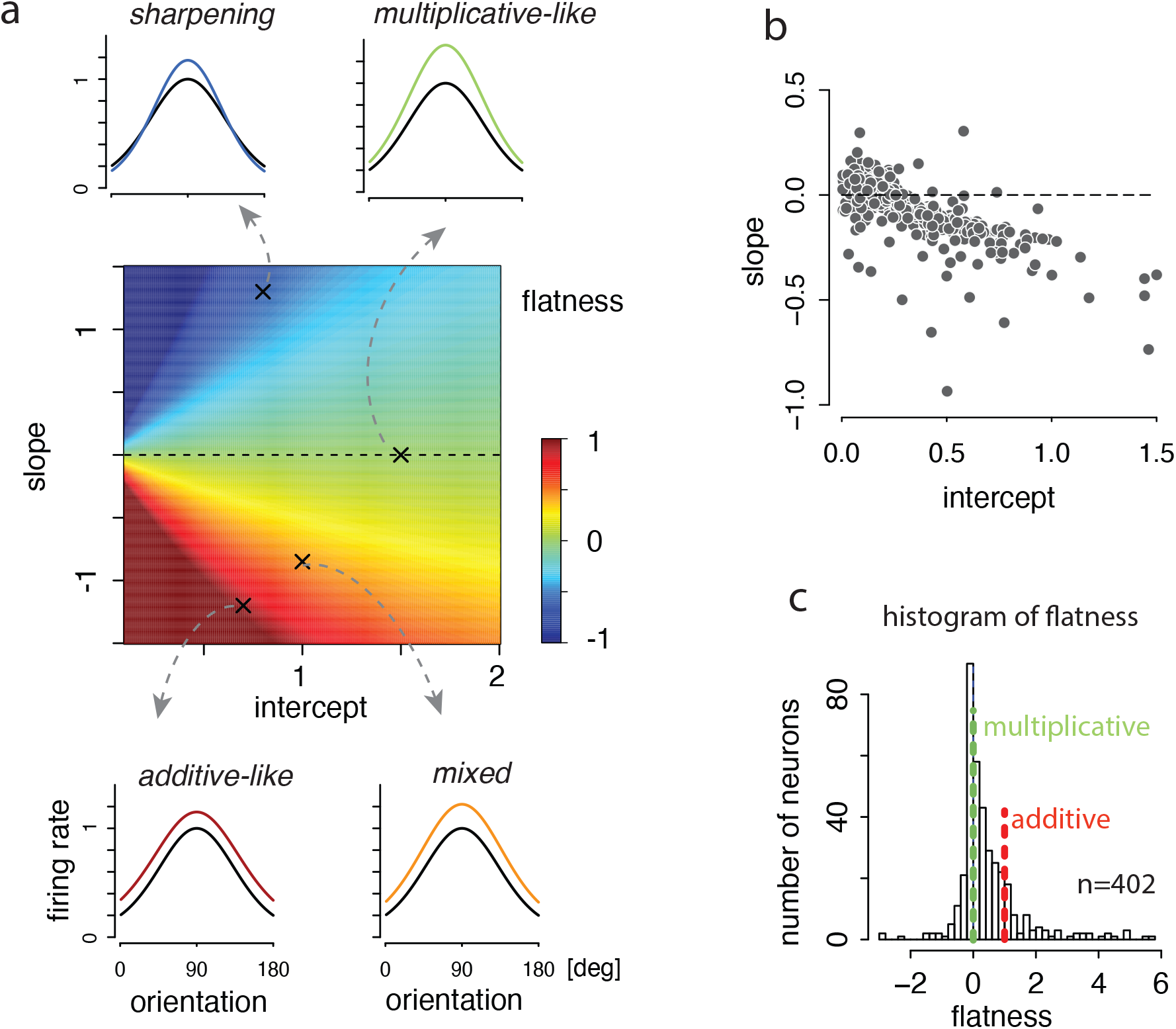
Power-law modulation accounts for both additive and multiplicative gain change. (a) The “flatness” index varies as a function of the slope and intercept parameters of the power-law relation. The “flatness” index quantifies the extent to which the firing rate change depends on the stimulus orientation. It manifests itself as multiplicative and additive gain, among others, in certain parameter regimes. Multiplicative gain would lead to a zero “flatness” index, while additive gain implies “flatness” equals to 1. Changing the flatness from 0 to 1 would lead to a gradual transition from multiplicative change to additive gain. This index can be smaller than 0, a case implying sharpening of the tuning curve. (b) The estimated values of slope and intercept for individual neurons. (c) The histogram for the flatness scores, indicating that tuning fluctuations lie on a continuum.

First, when the slope *w* = 0, the power-law modulation degenerates to a pure multiplicative gain [25]. Second, the power-law modulation could lead to approximately additive modulation with certain combinations of the slope(*w*) and intercept (*b*). For quantification, we define a “flatness” index to characterize the change over the stimulus variable induced by the fluctuation. Informally, this index computes the ratio between the change of the firing rates between the preferred and the orthogonal orientations (see Method for a formal definition). With additive gain, the flatness index is 1, while multiplicative gain leads to a flatness index of 0. When the flatness score is negative, the resulting configurations show sharpening of the tuning curve. Fig. 4 shows the “flatness” while systematically varying the two parameters (i.e., the slope and intercept). In the appropriate parameter regimes, the power-law would manifest itself as a multiplicative or an additive gain change (see Method for details), while parameter values in between resulting in tuning modulation which might be interpreted as a mixed of multiplicative and additive gains [28].

Empirically, most of the neurons lie in between multiplicative and additive gain change (Fig. 4b), thus are better characterized by the proposed power-law relation than a pure multiplicative or additive gain. The control analysis by generating simulated data using multiplicative gain showed that the recovered slope is close to zero over intercept as expected (Fig. S5). Note that [30] found that additive and multiplicative fluctuations were anti-correlated, which could be naturally explained by our power-law model. It is also worth mentioning that a subset of neurons exhibit a mild sharpening of the tuning curve. Together, these results provide a unified account of the fluctuations of orientation tuning in V1. Although we could not rule out the possibility of two separate mechanisms (one for multiplicative gain, and one for additive gain), our results show that a single form of fluctuation is sufficient to capture the variability, and the tuning fluctuations of individual neurons appear to lie on a continuum.

### 2.5 Population tuning fluctuations are low-dimensional

The dimensionality of tuning fluctuations has important implications of the mechanisms and the function of the circuit. Some studies (e.g., [28, 29]) implicitly assumed a rank-1 fluctuation that scales the gain of the population in a coherent manner, and found evidence suggesting that the total population activity is highly predictive of the moment-to-moment fluctuation of the response of individual neurons [29]. Others found the coupling strength of individual neurons to the rest of the local network to be diverse [55], implying a higher dimensionality of the tuning fluctuations and a potential role of recurrent connections in shaping network responses. Recently proposed E/I-balanced network models with spatial connectivity structure [56] predicted that the population fluctuations should be low-dimensional. Finally, a recent study proposed that gain variability in V1 serves to represent the stimulus uncertainty via sampling, a computation would generally require the gain variability to be high dimensional.

We examined the structure of the tuning fluctuation at the population level. As demonstrated earlier, for each neuron, the tuning fluctuations could be well captured by the first fPC. Exploiting this observation, we approximated tuning fluctuations of a neural population by concatenating the scores for individual neuron together (number of neurons × number of trials, Fig. 5a). Examining the correlation of the scores, we found that while most of the neurons fluctuate coherently, a small group of neurons are anti-correlated with the rest of the neurons (Fig. 5b) in some sessions (for results for all sessions, see Fig. S6). What is the dimensionality of latent fluctuations of the neural population? If the neurons share a coherent multiplicative gain change or additive gain change [28], the latent fluctuation should be close to one dimensional. To assess this, we performed a standard PCA analysis on the score matrix to assess the linear dimensionality. We found that, while the fluctuation shows low-dimensional structure (Fig. 5e), the dimensionality exceeds one.

**Figure 5:**
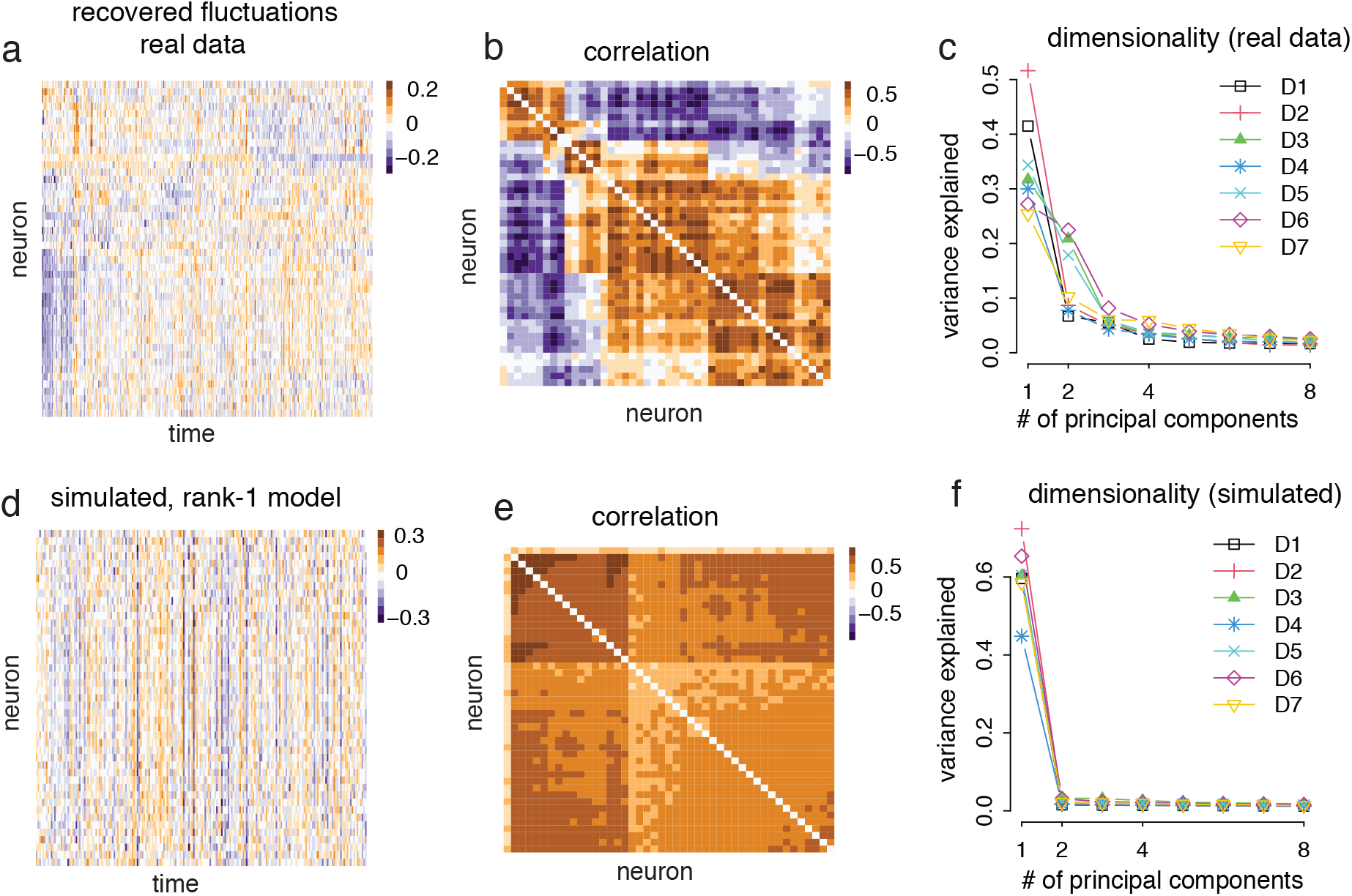
Population structure of the tuning fluctuations: the fluctuations of V1 population as captured by the first fPC have a low dimensional structure. (a) The heat map of scores of the first fPC for every neuron in one dataset (Session D5). Every data point represents the score of one neuron during one bock of stimulus presentation. (b) The correlation of the score matrix above. The neurons are sorted according to a hierarchical clustering algorithm in (a) and (b). (c) To quantify the dimensionality of the fluctuations as captured by the first fPC, we run a standard PCA on the score matrix. The first two principal components capture more than 60% of the variance. (d,e) Similar to (a,b), but based on the simulated dataset assuming rank-1 multiplicative gain fluctuations. (f) Similar to (c), but based on the simulated dataset assuming rank-1 multiplicative gain fluctuations. In this case, the first principle component consistently dominates other components in the recovered score matrix. These results suggest that the fluctuations in the neural population are low dimensional, but not one-dimensional.

The empirically observed latent fluctuations can not be explained by a rank-1 multiplicative or additive gain model. To demonstrate this, we performed control analysis by generating simulated data using the rank-1 additive gain or multiplicative gain model (see Method for details), and found that the resulting correlation structure of the inferred score based on the synthetic data exhibits simpler structure (Fig. 5d&e, and Fig. S7 & Fig. S8). The dimensionality of the scores is lower than that estimated from real data (Fig. 5f))

These results paint a more nuanced picture of the fluctuations of V1 at the neural population level. Deviating from what was suggested previously [28], the fluctuations of V1 neurons are not completely coherent in the anesthetized state, with subset of neurons could exhibit fluctuation at the opposite direction comparing to the majority, nor can it be characterized by a rank-1 additive or multiplicative gain fluctuations. These results will help further constrain and refine the network mechanisms giving rise to the tuning fluctuations in visual cortex [57, 56].

### 2.6 Higher neural activity barely increases, or even decreases Fisher information

So far, by applying our Pf-PCA analysis to the V1 data, we have derived a generative model of the neural activity in V1. Below, we demonstrate that this generative model is useful for characterizing various critical aspects of the neural code. In this section, we will leverage Pf-PCA to the calculation of the information to understand how the tuning fluctuation affects the information-carrying capacity of the V1 population. We focus on a local measure of the representation, i.e. Fisher information (FI), which has been important in quantifying the local property of the neural code [58, 59, 60, 61]. In the next section, we will use Pf-PCA to understand how the geometry of the neural response changes under tuning fluctuations, which represents another important aspect of the neural code. Overall, through the FI and geometry analysis, Pf-PCA enables several new understandings of the local and global structure of the V1 code.

First, using the model estimated by Pf-PCA, we examined the relationship between the FI and the magnitude of neural activity for individual neuron (Fig. 6a,b). See Method for the calculation of the FI. We found that this relationship differs substantially from neuron to neuron- it could be positive, negative, or flat (Fig. 6a,b). Fig. 6b shows the histogram of the slope when regressing FI against the neural activity. Interestingly, the median of these slopes is close to 0 (i.e., 0.001). We further validated these results by performing recovery analysis using synthetic data. We found that our method could indeed faithfully recover the relationship between FI and neural activity for individual neurons given the particular sample size of the data (see Fig. S9).

**Figure 6:**
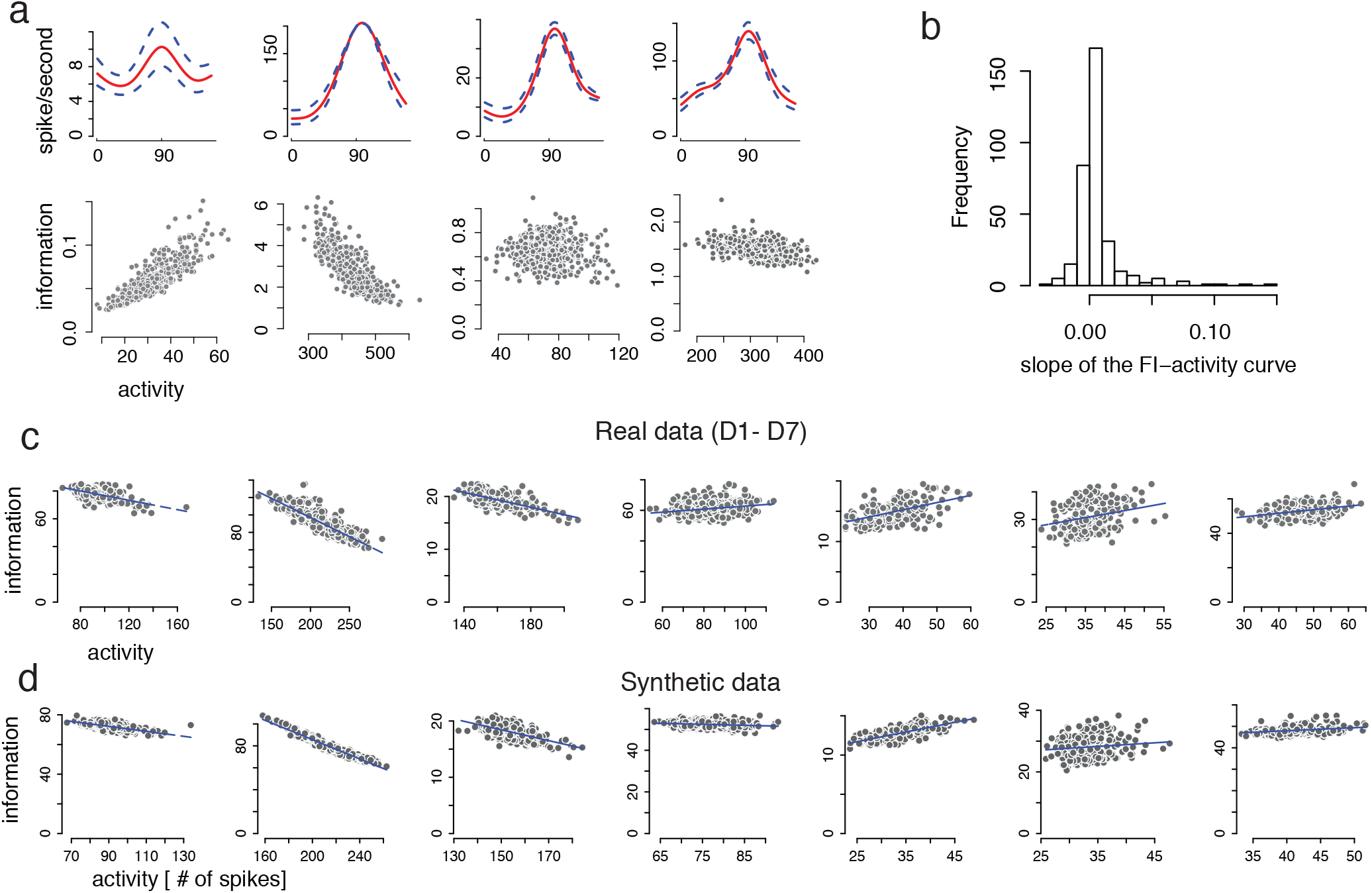
Fisher information analysis reveals that, higher neural activity barely increases, or even decreases Fisher information. (a) The relation between the FI and the neural activity (lower panels) and the tuning curves with latent fluctuation induced by the first fPC (upper panels) in four neuron examples. (b) Histogram of slopes of the FI-activity for all neurons (n=402). (c) Scatter plots showing the relationship between the population FI and the neural activity for each session in real data. (d) Scatter plots showing the relationship between the population FI and the neural activity for the synthetic data. The synthetic data were generated from the Pf-PCA model fit to each corresponding dataset. Note that there is a tendency for the model to under-estimate the scores (as expected), leading to a proportional under-estimation of the population FI. Importantly, this under-estimation does not affect the recovery of the relation between the population FI and the neural activity.

Fig. 6c shows the population FI for all neurons in each dataset, sorted according to the neural activity. Here we assume that the neurons are noise-independent conditioned on the latent fluctuations. Note that a multiplicative gain model predicts that the population FI scales proportionally with the amount of neural activity, or put it in another way, doubling the firing rate would doubling the population FI. However, we found that, for most sessions, the population FI is minimally affected, or decreases systematically as the neural activity increases. This is in sharp contrast with the multiplicative gain model. To quantify this, we defined a FI-modulation index (i.e., the slope of FI-activity curve). With multiplicative gain model, this FI-modulation is exactly 1. In the data, the modulation indexes for all sessions are far smaller than 1 and in some cases negative (−0.17, -0.44, -0.07, 0.10, 0.12, 0.26, and 0.21, respectively). Recovery analysis based on synthetic data suggests that our procedure could indeed recover the relationship between the neural activity and the population FI (Fig. 6d). See Method for details. These results are consistent with [30] in that both studies found that increased neural activities do not lead to substantial increase of the population FI. Meanwhile, we also noticed some subtle discrepancies between the two studies, because [30] suggested that there was minimal change of the population FI when neural activity changes. We believe that the difference lies in the difference in the analysis methods (see Method and SI.6 for a detailed discussion).

### 2.7 Change of representational geometry induced by spontaneous fluctuation is different from contrast

Neural response naturally forms neural manifold by systematically varying the stimulus. The geometry of the encoding manifold has multiple implications in understanding the format of the representation and in linking neural responses to the behavior (reviewed in [62]). To understand how the tuning variability affects the geometrical properties of the encoding manifold, we begin by simulating a multiplicative gain change, which is simple scenario exhibiting how fluctuating signals are displayed by the geometric analysis. We constructed a homogeneous population code for encoding stimulus orientation with independent Poisson noise, and varying the shared multiplicative gain (see Method for details). This population coding model recapitulates the basic effect of varying stimulus contrast [22, 43, 23]. We computed the representational distance [63] as a function of the orientation disparity, and found that multiplicative gain only scales the representational distance function without changing the shape (Fig. 7c). A 3-D multi-dimensional scaling (MDS) based on the representational distance matrix shows that the neural manifold under tuning fluctuation exhibits a cone-shape (Fig. 7d), with the radial dimension encoding the multiplicative gain. When projecting onto the first two dimensions, we observed that the size of the representation for each contrast (e.g., the radius of each circle) scales with the neural activity (Fig. 7e).

**Figure 7:**
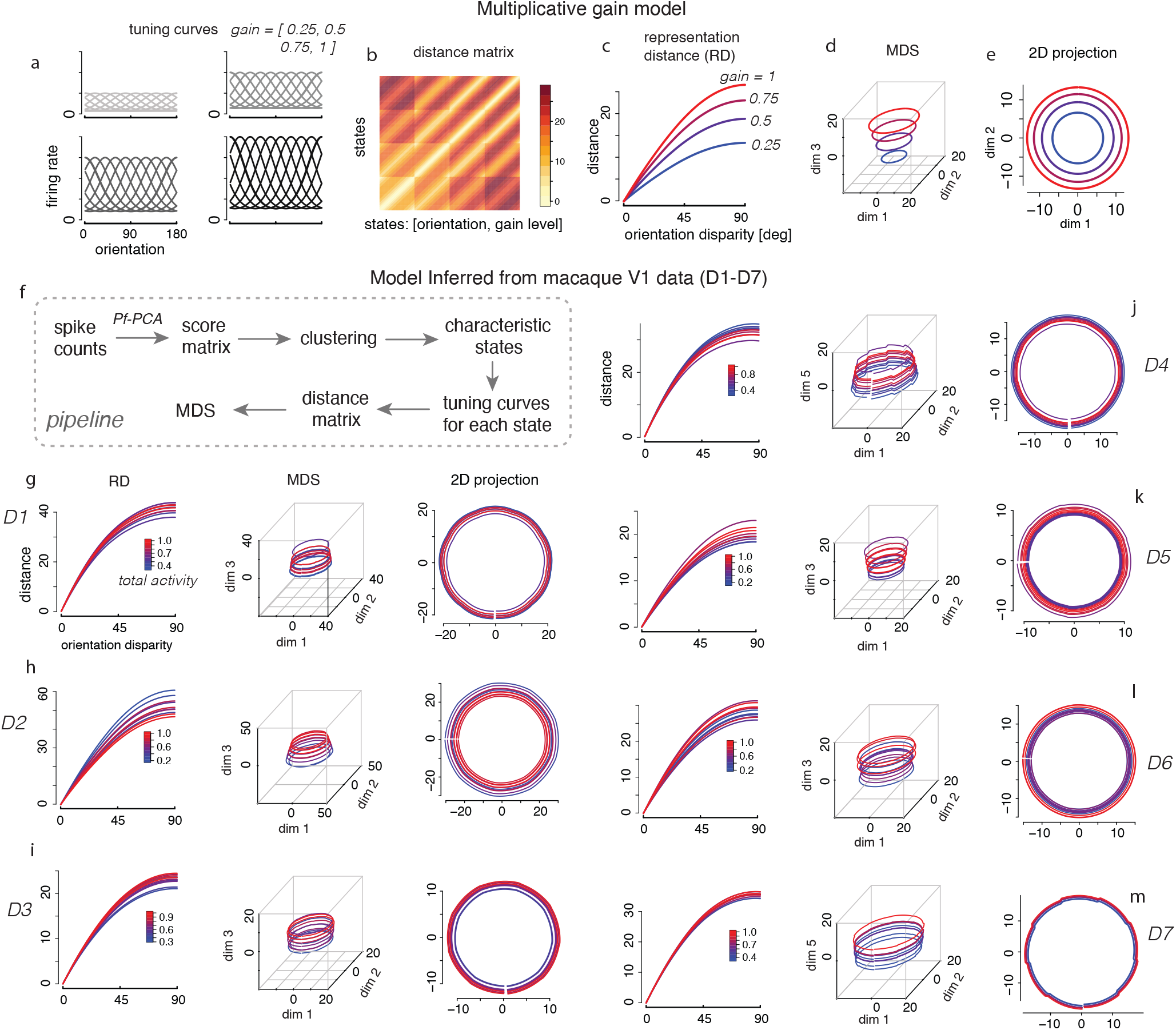
Geometry analysis demonstrates that the latent tuning fluctuations lie on a different manifold comparing to changing contrast. (a-e) Analysis based on multiplicative gain model. (a) Tuning curves under different multiplicative gain (or contrast) levels. (b) The representational distance matrix for each pair of states, defined by the orientation and the gain level. We discretized the orientation into 180 bins, results in 180× 4 = 720 states. The states were arranged according to the orientation and states. (c) The representational distance (RD) as a function of orientation disparity for four different gain levels. (d) 3-D MDS reveals a cone-like structure of the neural population code. (e) Projection onto the first two dimensions reveals that the size of the representation (measured by radius) scales proportionally with the neural activity. (f) Pipeline of geometric analysis based on real data. For each session, we inferred the score matrix for the first fPC, and then computed ten characteristic states using a clustering procedure. For each characteristic state, we generated the corresponding tuning curves using the results from Pf-PCA. The curves for different states are color-coded based on the total activity (normalized to the maximum). (g-m) The results for each dataset from the geometric analysis. The MDS results reveal cylinder-like structure in most of the sessions. Although total activity can change substantially across the different scales, however, the size of the representation does not change substantially. This is apparent when comparing to the multiplicative gain model.

Next, we sought to understand how the tuning fluctuations identified by Pf-PCA from the V1 data would affect the geometry of the code [64, 65]. To do so, we created a neural population code based on the empirically-fitted tuning curves and scores from Pf-PCA (see Method for details). We clustered the score matrix into 10 clusters, then computed the average scores for each cluster to get the pattern of fluctuations corresponding to each of the 10 characteristic states. For each state, the corresponding tuning curves were generated accordingly. Analyzing the representational distance (RD) as a function of orientation disparity for the 10 latent states (shown in Fig. 7g-m, first column), we found that the RD curves are only slightly affected by the total activity. In several cases, higher activity leads to overall lower RD, e.g., (Fig. 7h). Furthermore, MDS analysis (Fig. 7g-m, second column) shows that the fluctuations cause the representation to move along a cylinder-like manifold. When projecting onto the first two dimensions, we had two observations. First, the centers of the representations corresponding to each states are aligned, suggesting that the representation “drifts” [66] in the direction that is orthogonal to the representation of orientation. Second, the size of the representation only changes slightly with varying population activities (Fig. 7g-m, third column). Note that this general pattern does not resemble the cone-like structure induced by the multiplicative gain (Fig. 7d). Note that in two of seven sessions, the latent fluctuations are smaller so that the cylinder structure does not appear in the 3-D MDS, however, it becomes apparent when we plotted the first two and the fifth dimension in a 5-D MDS embedding.

These results demonstrate that the spontaneous fluctuations of neural tuning in V1 lie on a different manifold compared to that induced by the changing stimulus contrast that leads to a multiplicative gain. Note that the effect is also different from a simple additive gain (Fig. S11). These results have important implications for the downstream readout. If the spontaneous fluctuations lied on the same manifold as the changing stimulus contrast, the downstream would not be possible to distinguish the spontaneous fluctuations from a change of contrast. Our results argue against that scenario, and further suggest that the latent fluctuations mostly cause a “drift” of the representation without fundamentally changing the fidelity and the structure of the representation [66, 67, 68].

## 3 Discussion

We have presented a flexible unsupervised approach, Pf-PCA, to analyze the tuning variability. This approach provides a general framework to understand how the observed stimulus variables and latent factors together influence the neural activity. Specifically, it decomposes the tuning curves as the sum of the mean component and the fPCs which are tuned to the stimulus, and are subject to the modulation of the latent factors. We demonstrated that Pf-PCA could robustly and reliably recover the structure of the fluctuations given a few dozens of trials of data. We applied our new method to analyze the spike train data collected from anesthetized macaque V1 while viewing drift gratings, and discovered several novel insights regarding the structure of the orientation code.

Our method represents a more flexible modeling framework compared to previous work in analyzing the tuning variability. Previous models often presumed the form of fluctuation (e.g. [25, 28, 30, 31]), and the fluctuation was often assumed to be a constant, acting onto the tuning curve through either multiplicative or additive interactions. Thus, the forms of fluctuations captured by these analyses were limited by construction. Our method, instead, allows unsupervised discovery of arbitrary smooth tuning fluctuation, and potentially multiple forms of fluctuations simultaneously. Our method is broadly applicable, so long as the neurons have smooth tuning over certain stimulus dimension, which could be spatial frequency [69], location [70, 17], direction [71, 72], or time [73]. A potentially fruitful venue of using our approach would be to leverage our approach to testing computational models by analyzing the data simulated from these models to identify the structure of the latent fluctuations, and comparing these predictions with the structure extracted from the data.

We have focused on an exponential non-linearity for the link function, which has been assumed by many previous models [74, 75, 76, 77]. It should be possible to further extend it to other types of nonlinearity [78], such as a power-law transformation [43, 79]. It would also be interesting for future research to develop techniques that could automatically infer the type of non-linearity from the data directly.

Our V1 results should be informative to a better mechanistic and functional understanding of the V1. Naively, assuming an exponential non-linearity, the power-law modulation revealed by our analysis could be explained by a tuned input to a given neuron which fluctuates over time. However, this is likely a simplified picture. It would be more fruitful to consider how threshold non-linearity together with noise could lead to these kind of results. Previously, models of this kind [79, 80, 81] have been used to account for the multiplicative gain on the tuning curve induced by varying contrast. Second, the finding that the latent fluctuations are heterogeneous in the population is consistent with the idea that the recurrent processing in V1 may play an important rule in shaping the structure of the fluctuations of neural tuning. These results echo with recent work [56] showing that spatially patterned fluctuation structure could emerge in balance networks in V1 in which neural fluctuations can be heterogeneous. Third, it is interesting to consider the implications of our observations in the context of the functional models of neural variability. Such variability has been proposed to reflect sampling of the sensory inputs [82], encoding stimulus uncertainty [27], and efficient encoding of natural scene statistics [83, 84, 85]. The specific structure of the latent fluctuations extracted by Pf-PCA provides a richer set of summary statistics to further test these current mechanistic and functional models and help developing future models.

Our method enables us to further analyze the coding properties in the presence of the tuning fluctuations, both in terms of local properties (via FI) and the global geometrical structure of the code under tuning fluctuations. We found that FI generally does not substantially increase (sometimes even decrease) with increased neural activity. This may point to the potential importance of cortical inhibition in sharpening the neural code [86]. The analysis of the geometry reveals that the manifold induced by the latent fluctuation lies in different subspace of changing contrast. This suggests that the tuning fluctuations in V1 may not interfere with encoding of contrast. These observations deserve further investigations in future.

A few limitations and the potential improvement of our results are worth mentioning. First, our V1 results are based on analyzing neural responses at the anesthetized state. It would be interesting to examine whether tuning fluctuations of awake behaving animals would follow similar rules. It would be interesting to see if the fluctuations of internal states correspond to a change of the behavior [68]. Also, it should be possible to improve our method by leveraging temporal smoothness prior on the scores, e.g., by assuming a Gaussian process prior [87, 88, 89, 90, 91, 92]- a direction we did not pursue here, but would be an interesting future direction. Another limitation is that our method when applying to V1 data only deals with slow fluctuations [25], because of the assumption that the latent is the same across every block (*∼* 10 seconds). Thus our estimate of the magnitude of the latent fluctuation is likely an under-estimate of the true fluctuations. Methodologically, these issues may be addressed in two ways: by using faster stimulus sampling in experiments; ii) by developing methods to fit the neural population all together. For the latter, assuming a low dimensional latent structure, it should be possible to infer the latent fluctuation based on individual stimulus- a direction we are currently pursuing.

In summary, we have developed a novel statistical approach to parse the variability of neural tuning. Our approach can flexibly capture the impact of both stimulus variable and latent variable on a trial-by-trial basis. Applying our approach to macaque V1 revealed the structure of the tuning variability both at the level of individual neuron and the neural population. Our analyses also led to further insights on the FI and geometry of the code. While we only analyzed the orientation code for V1 in the paper, we hope that the analysis pipeline developed here would be informative for elucidating the structure of the tuning in other neural systems as well [93].

## Acknowledgement

We thank Adam Kohn, Matt Smith, and Inigo Arandia-Romero for sharing the V1 data. We thank Liam Paninski, Robbe Goris and Nikolaus Kriegeskorte for fruitful discussions. We thank Kenneth Kay, Yvonne Li, Matthew Whiteway, and Mattia Rigotti for comments on an earlier version of this paper.

## 4 Methods

### 4.1 Poisson functional principal component analysis (Pf-PCA): generative model

A standard model description of neural responses in neuroscience is based on tuning curves and (typically) Poisson spiking noise. Specifically, the observed spike count *n*(*s*) of stimulus *s* for individual neuron, during a counting window of length Δ_*t*_, is modelled as a Poisson distribution

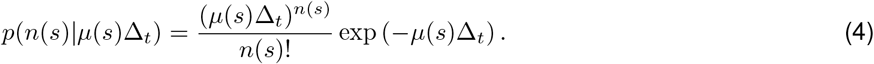

where *µ*(*s*) represents the tuning curve of stimulus *s*.

The tuning curve may vary among *B* trials and is not directly observed. Denoting the tuning curve for each trial as *µ*_*t*_(*s*) for trial *t, t* = 1, … *B*, we model the log of the stochastic curve, log(*µ*_*t*_(*s*)), as following:

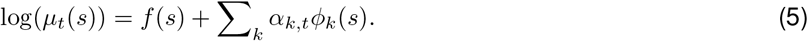

Here *f* (*s*) is the *mean component, φ*_*k*_(*s*) is the *k*-th functional principal component (fPC), and *α*_*k,t*_ denotes the amount of fluctuation (i.e., score) for the *k*-th component during the *t*-th trial, and is assumed to follow a zero-mean Gaussian distribution, with variance 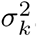. This model implies that log(*µ*_*t*_(*s*)) has the mean E[log(*µ*_*t*_(*s*))] = *f* (*s*) and the co-variance function 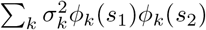.

Note that with only the first term *f* (*s*), this model is equivalent to the standard tuning curve model of spike counts. The second term Σ_*k*_ *α*_*k,t*_*ϕ*_*k*_(*s*) is the additional variance from the contribution of trial-to-trial fluctuations, making the model naturally capture the over-dispersion of spike counts.

Pf-PCA differs from the multiplicative gain model proposed in [25]. First and most importantly, we do not assume a pure multiplicative gain change in the fluctuations, instead, the form of fluctuations is arbitrary. Second and a more subtle point is that, in Pf-PCA, the magnitude of the fluctuation *α* are assumed to follow a Gaussian distribution, while Gamma distribution is assumed for the gain in [25] in the firing rate scale (not the logarithm of firing rate). Notice that *α*_1,*t*_ is Gaussian distributed thus symmetric, while the logarithm of Gamma distribution is left-skewed thus has a different shape.

### 4.2 Inference: a two-step estimation procedure

Assume that we have observations based on a set of *m* stimuli which are sampled from a particular stimulus space *S*. The spike count of individual neuron elicited by individual stimulus *s*_*j*_, *j* = 1, …, *m*, is denoted as *n*_*t*_(*s*_*j*_) for the *t*-th trial. We further denote the spike count vector for the *t*-th trial as 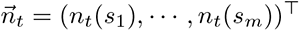.

Intuitively, if we could recover the unobservable mean *µ*_*t*_(*s*_*j*_) for *t* = 1,, *B, j* = 1,, *m*, fitting the model of stochastic curve log(*µ*_*t*_(*s*)) using functional PCA would be straight-forward. Denote 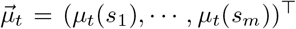. We could estimate the posterior of the hidden firing rate 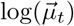 from the spike count data using an expectation-maximization (EM) algorithm [94]. Following these ideas, we developed a two-step estimation procedure, as following.

#### Step 1: recover the hidden 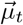

When the vector 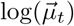 is observable, the likelihood of Poisson model can be written as

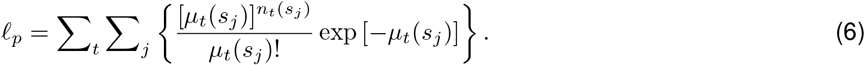

Eqn. (5) implies that the logarithm of the firing rate for the sampled stimulus set {*s*_1_, …, *s*_*m*_} during the *t*-th trial 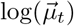 can be modeled as 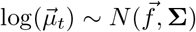, where 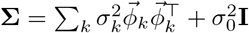. We then obtain the log-likelihood

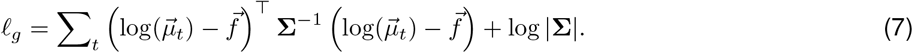

Although in reality the firing rates are not directly observed, however, by treating 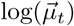 as missing data, we could use an EM algorithm to iterate between an E-step and a M-step to optimize the functions. Specifically, the E-step calculates the conditional mean 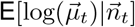 and conditional variance 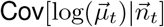, given the current estimates of the parameters 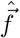 and 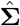 obtained in M-step. The M-step maximizes the first term of Eqn. (7), which involves the following two quantities:

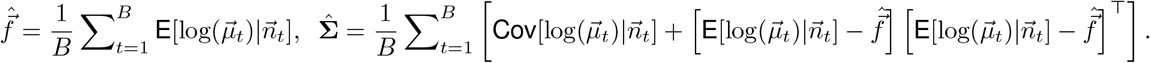

Note that calculating these expectations in the E-step requires the marginal distribution 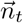, which is not analytically tractable. We thus adopt a Monte Carlo approach to calculate them. Together, **Step 1** gives an estimator of hidden means 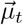 in the form of 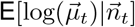.

#### Step 2: perform functional PCA on the recovered hidden 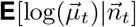

Given the posterior 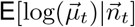, we then apply the methods of functional data analysis into the estimated posterior to get the mean component *f* (*s*), the fPCs *ϕ*_*k*_(*s*), and their corresponding scores *α*_*k,t*_. Specifically, *f* (*s*) is obtained by using the natural cubic splines smoothing approach

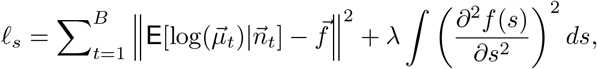

where *λ* is chosen via generalized cross validation [95].

The functional fluctuations { *ϕ*_*k*_(*s*)} are estimated by the roughness of the eigenfunction [96, 51]. The first component 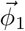 and the corresponding score for each trial 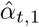 are estimated via

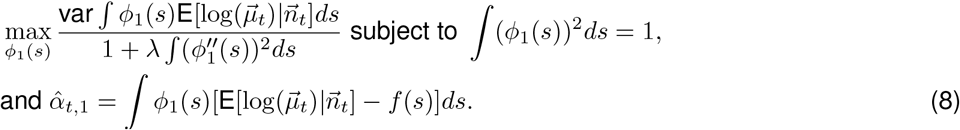

The remaining components and their scores are obtained via an iterative process such that any higher order eigenfunction is orthogonal to the eigenfunctions already recovered. This procedure allows us to estimate the variance explained by each fPC, as quantified by the variance of the score for that component. The proportion of variance explained by the *k*-th fPC 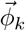 is simply calculated by 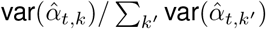.

### 4.3 Implementation of a reduced version of the method: *µ*-PCA

We also implement a reduced version of the method, *µ*-PCA. For this method, after obtaining the posterior mean 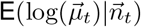, we perform regular PCA directly on the exponential function of the estimated posterior mean, 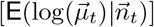, instead of functional PCA,

*µ*-PCA can be thought as an alternative way to perform PCA to Poisson count data compared to Poisson PCA [53]. First, [53] considered log(*µ*_*t*_(*s*)) as “natural parameters”, which define the mean and components. In contrast, *µ*-PCA considers the unobserved log(*µ*_*t*_(*s*)) as random variables, and decomposes it into the mean and the principal components with the amount of fluctuations assumed to be Gaussian. Second, different techniques are proposed to obtain the principal components. Following the relation between log-likelihood of exponential family and Bregman distance, [53] constructed a loss function to optimize it, then obtained the principal components. The *µ*-PCA method seeks to estimate the posterior mean 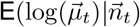 instead of random observable *µ*_*t*_(*s*).

### 4.4 Validation of the methods using simulated data

We validated our methods using simulated data with ground truth. To do so, we generated tuning curve 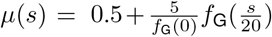 where *f*_G_ is the density function of standard normal distribution. The stimulus *s* takes a sequence with a 22.5 degree interval from -90 to 90 degree. The number of trials *B* was set to be 50 throughout.

For each of the four types of tuning changes, i.e., multiplicative gain, additive gain, tuning shift, and tuning sharpening, we reverse-engineered the fPC 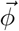 that would give rise to that type of tuning fluctuations. More concretely, the tuning fluctuations, denoted by 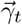, are generated from Gaussian with covariance structure 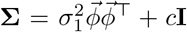, where *c* is chosen such that the structured component explains 80% of the variance. The second component amounts to white noise, accounting for 20% of the variance. Adding this random noise component allows us to test the robustness of the model in the presence of less structured fluctuations, as well as to evaluate the extent to which our estimation procedure could recover the variance explained by the structured fPC. Note that the recovery problem becomes easier without such random noise component. From the simulated data, we first estimated the variance explained by the *K* fPCs (*K* = number of stimuli) using Pf-PCA, and then computed the proportion of variance explained by the first fPC as 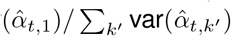. Effectively, we relied on the last *K* − 1 components to recover the random component in the generative model.

Here *µ*_0_(*s*) = exp(*f* (*s*)) represents the tuning curve with no fluctuation, and 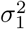 denotes the the strength of fluctuation. Due to the scale difference among different fluctuations, the parameter 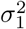 is set as following. Multiplicative gain: 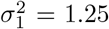, so that its gain change is log(1.3*µ*_0_(*s*)) *−* log(0.9*µ*_0_(*s*)). Additive gain: 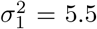, so that its gain change is log[*µ*_0_(*s*) + 0.4] − log[*µ*_0_(*s*) − 0.2]. Tuning shift: 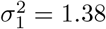, so that it shifts log(*µ*_0_(*s* + 6)) − log(*µ*_0_(*s −* 6)). Tuning width change: 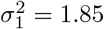, so that its standard deviance changes ±20%. Under these settings, we generated Poisson spike count data with firing rates *µ*_*t*_(*s*) = exp[log(*µ*_0_(*s*)) + *γ*_*t*_(*s*)].

Note that we also validated our methods i) when multiple forms of fluctuations co-exist simultaneously; ii) when the tuning curves are monotonic. These procedures and their results can be found in SI (see SI.1 & SI.2).

### 4.5 Analyzing data from macaque V1 using Pf-PCA

We used 7 datasets which were published previously. Three of them (D5-D7), publicly available from CRCNS website, were obtained from anesthetized macaque primary visual cortex by Matthew Smith and Adam Kohn [54]. In these experiments, described in details in [54, 97], spiking activities were recorded while presenting different grayscale visual stimuli, including drifting sinusoidal gratings (each presented for 1.28 s). These gratings are [0, 30, 60, …, 330] deg. The other four sessions (D1-D4) were previously published in [30], shared by the authors. These data also visually evoked activities from anesthetized macaque primary visual cortex (see [30] for details). The grating directions are [0, 22.5, 45, …, 157.5] deg. We chose neurons with SNR ≥ 2 and mean firing rate ≥ 1.5 spikes/second. In total, we analyzed 7 datasets with 402 neurons.

For each stimulus, we counted the number of spikes for a 500ms window (80ms - 580ms after stimulus onset). Because the experiments had a block-randomized design, for each block we obtained a response vector corresponding the responses for all the stimulus orientations sampled in the experiments. Repeating this for every block, we constructed a spike count matrix for each neuron (number of trials × number of orientations).

We then applied Pf-PCA to this matrix for each neuron. In doing so, we obtained the mean component *f* (*s*), the fPCs *ϕ*_*k*_(*s*), where *k* is component index, as well as the amount of fluctuations, i.e., scores, *α*_*k,t*_ for each trial *t*. We set the number of fPCs to be to three, as three fPCs could already sufficient to account for most of the variance (see Fig. S4).

In Fig.3a, we plotted the inferred mean component in the form of exp(*f* (*s*)), and the first fPC in the form of exp(*f* (*s*) *± σϕ*_1_(*s*)), where *σ* is the s.d. of estimated *α*_1,*t*_.

#### Regression analysis

we analyzed the relationship between *ϕ*_1_(*s*) and *f* (*s*) by performing a regression analysis with the following form

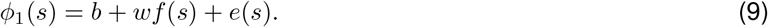

The regression is done by using the “lm” function in scientific computing software R. First, we tested the significance of the regression by a F-test (see Fig. 3c). To quantify how much information of *ϕ*_1_(*s*) is accounted for by a linear function of *f* (*s*), we defined a summary statistics: 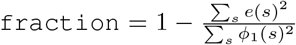. This measure was reported in Fig. 3d.

#### “Flatness index” analysis

In Fig. 4a, *µ*_0_(*s*) = exp(*f* (*s*)) was generated from von Mises function with parameters satisfying *µ*_0_(*s*) ∈ [0.2, 1]. Denote the tuning curve corresponding to the fluctuation *α* = *α*_0_ as 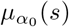. Define Δ*µ*(*s*) = *µ*_*α*_(*s*) − *µ*_0_(*s*) − *c*(exp(*bα*) − 1), where *c* is the baseline of *µ*_0_(*s*). Thus, Δ*µ*(*s*) captures the change of firing rate as a function of the stimulus with an additional correction term. The “`flatness`” index was defined as 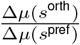, where *s*^pref^ and *s*^orth^ denote the preferred orientation of the neuron and its orthogonal orientation, respectively.

This index quantifies how flat Δ*µ*(*s*) is. For additive gain change, *µ*_*α*_(*s*) = *µ*_0_(*s*) + *c*_add_*α*, where *c*_add_ is a constant, and Δ*µ*(*s*) = *c*_add_*α* − *c*(exp(*bα*) − 1), implying Δ*µ*(*s*) is completely flat over *s*. Thus, `flatness` = 1 in this case. For multiplicative gain change, *µ*_*α*_(*s*) = exp(*bα*)*µ*_0_(*s*) and Δ*µ*(*s*) = (*µ*_0_(*s*) − *c*)(exp(*bα*) − 1), implying Δ*µ*(*s*^orth^) = 0. Thus, `flatness` = 0. Remarks: `flatness` can be larger than 1 when Δ*µ*(*s*^orth^) *>* Δ*µ*(*s*^pref^), which is possible when Δ*µ*(*s*) *<* 0. It is also possible to have `flatness` *<* 0, when Δ*µ*(*s*^orth^) *<* 0 *<* Δ*µ*(*s*^pref^).

#### Connecting the power-law relation to multiplicative gain and additive gain

The power-law modulation can degenerate to multiplicative gain and additive gain under certain parameter regime. Obviously, as *w* = 0, the power-law modulation is equivalent to multiplicative gain. The connection to additive gain is less obvious. When *bα* and *wα* are close to 0, using Taylor expansion, *µ*_*α*_(*s*) ≈ *µ*_0_(*s*) + *µ*_0_(*s*)(*b* + *w* log *µ*_0_(*s*))*α*. It follows that when the function *µ*_0_(*s*)*b/w* +*µ*_0_(*s*) log *µ*_0_(*s*) is flat over *s*, the power-law modulation degenerates to additive gain. Examining the property of the function *g*(*x*) = *x* log(*x*) − *κx*, we found that *g*(*x*) is indeed approximately flat over [0, 180] when *κ* is in some region, resulting in an approximate additive gain modulation.

### 4.6 Control analysis

#### Simulated data from multiplicative gain model

To generate synthetic data from the multiplicative gain model, we set the tuning curve *µ*(*s*) on each trial to be log(mean firing rate) plus a constant fluctuation, with its standard deviation matching that was inferred from the real data. We then sampled the spike count under Poisson noise. In doing so, we generated synthetic data which approximately match the amount of fluctuations in the real data, but with a pure multiplicative gain change. We performed regression analysis on the simulated data using the same procedure as that was used for real data (described above). The slope values obtained for the real and synthetic data were reported and compared in Fig. 3e.

#### Rank-1 model

In Fig 5c,d,f and SI Fig S7, we reported the recovered score matrix based on a Rank-1 multiplicative-gain model and the dimensionality. When simulating this model to generate synthetic data, the multiplicative gain fluctuations were sampled i.i.d. from a normal distribution. The fluctuation in each trial was shared by all neurons in the population, ensuring the score matrix is rank-1. We performed the Pf-PCA analysis on the simulated population, then obtained the recovered score matrix. We also simulated and analyzed synthetic data from a rank-1 additive-gain model. These results were reported in SI Fig S8.

### 4.7 Fisher information

Assume that tuning curve for neuron *i* is *µ*^(*i*)^(*s*) and that the spike count *n*^(*i*)^(*s*) follows Poisson distribution with mean *µ*^(*i*)^(*s*). Because we can approximate *µ*^(*i*)^(*s*) by the mean *f* (*s*) plus the functional fluctuations, the Fisher information (FI) of neuron *i* at stimulus *s*, given the score *α*_*k*_, where *k* is index of component, is obtained by

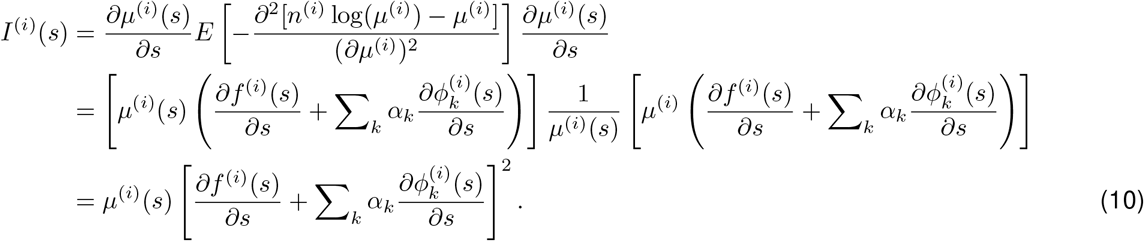

To compute the population Fisher information, we assumed that the neurons are independent conditioned on the fluctuations. To compute the population FI for each stimulus, we summed over the neurons in the population. Note that the reported FI (for individual neuron or neural population) is the total FI by summing over different stimulus orientations.

#### Recovery analysis on FI

To see if our method indeed has the statistical accuracy to recover the relation between FI and spike activity, we performed a control recovery analysis. We first generated synthetic datasets by simulating data based on the Pf-PCA model with the parameter values estimated from the real data. Specifically, for a neuron we considered the Poisson mean *µ*_*t*_(*s*) of trial *t* to be log(*µ*_*t*_(*s*)) = *f* (*s*) + *α*_1,*t*_*ϕ*_1_(*s*), and generated the counts of trial *t* from Poisson with the mean. From this, we obtained the synthetic population counts. We then performed the same analysis pipeline on these synthetic data to estimate the population FI. From this control analysis, we found our method can accurately recover the relationship between FI and total spiking activity.

#### FI & classification analysis

We performed classification analysis similar to [30] to examine the relation between the population FI and classification accuracy. Similar to [30], we splitted the data into two groups (i.e., high and low), sorted by the population activity. We performed classification based on ensemble with different size. Given a randomly selected ensemble of neurons with certain size, we performed multinomial logistic regression, and obtained the performance (proportion of correct classification). For avoiding over fitting the data, we used 5-fold cross-validation and reported the average performance across the five sets of left-out data. For each ensemble size, we performed this analysis on 500 randomly selective groups for high and low group each. These results were reported in Fig. S10(a) in the SI. We also performed this analysis on the synthetic data as described above. The results were shown in Fig. S10(b).

### 4.8 Analysis of representational geometry

We analyzed the geometry of the representation under a simple multiplicative gain model and the power-law model infer from the V1 data.

For the simple multiplicative gain model recapitulating the effect of changing contrast, we generated a homogeneous set of tuning curves using von Mises function (tuning width parameter equals 1). We assumed that the multiplicative gain modulated the firing rate of all neurons in the same way. In Fig. 7a-e, we assumed that the multiplicative gain could take four different levels (0.25,0.5,0.75,1), and computed the representation distance matrix by evaluating the representational distance for each pair of states (defined by both stimulus orientation and multiplicative gain). We performed 3-dimensional classic MDS to visualize the geometrical structure of the representation, and obtained the projection onto the first two dimensions. A similar analysis was performed based on pure additive gain change (for results, see Fig. S11).

For the models based on Pf-PCA inferred from real data (Fig. 7f-l), we performed the geometry analysis by the following steps:

i. We first generated the mean firing rates for each trial *t* by from our power-law modulation model. We clustered the trial × neuron score matrix into 10 clusters according to trials by k-mean, then computed the average score within clusters to get the “10-state averaged score”, which is 10 × neuron matrix. For each of the 10 states in the population, the corresponding tuning curves were generated.
ii. To reduce the biased sampling of neurons, we created a more shift-invariant neural population code by shifting tuning curves 8 times with 20 degree each. Our assumption here is that the neural code for orientation in V1 is roughly shift-invariant.
iii. We calculated the euclidean distances between stimuli based on the extended population matrix (after variance-stabilizing transformation for Poisson noise, i.e., taking the square root transformation) to obtain a distance matrix, and performed the classic MDS based on this distance matrix. In most of the sessions, we performed 3-D MDS. In two of seven sessions, the latent fluctuations are smaller so that the cylinder structure does not appear in 3-D MDS. For these two sessions, we performed 5-D MDS. When plotting the first two and the fifth dimension in a 5-D MDS embedding, the cylinder-like structure is apparent.

## Supplementary information

### SI.1 Recovering mixed fluctuations

Fig. 2 and Fig. S3 have shown that the tuning fluctuations could be captured by a single functional component, however, fluctuations in real data may consist of a mixture of different kinds of variability, e.g., lateral shift and the multiplicative gain. Therefore we next examine if our method would work given such more complex structure. We mixed two of the single kinds above (see Fig. S1) and stimulated the count matrices from the neural tuning. Fig. S1 shows that the recovered components from Pf-PCA are close to the ground-truth fluctuation, which suggest that the method can also successfully identify latent structure in more complex cases (see Fig. S1).

**Figure S1:**
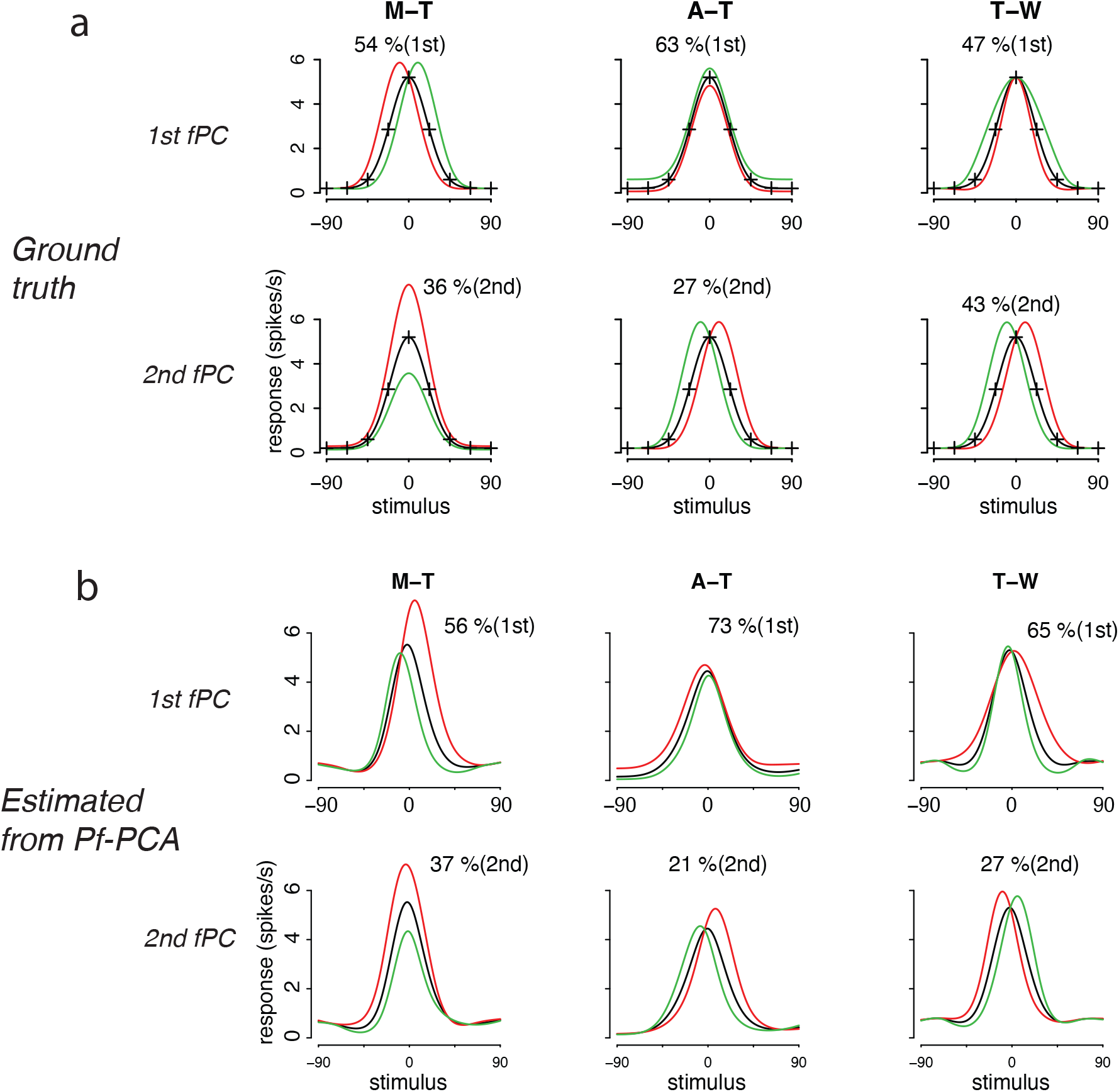
Recovering mixed fluctuations with two components(related to Fig. 2). (a) Ground truth. M: multiplicative gain; T: tuning shift; W: tuning width change. M-T represents mixing multiplicative gain and tuning shift; same convention for the other two cases. (b) Components recovered by Pf-PCA. The percentages show the proportions of variance explained by individual component.

**Figure S2:**
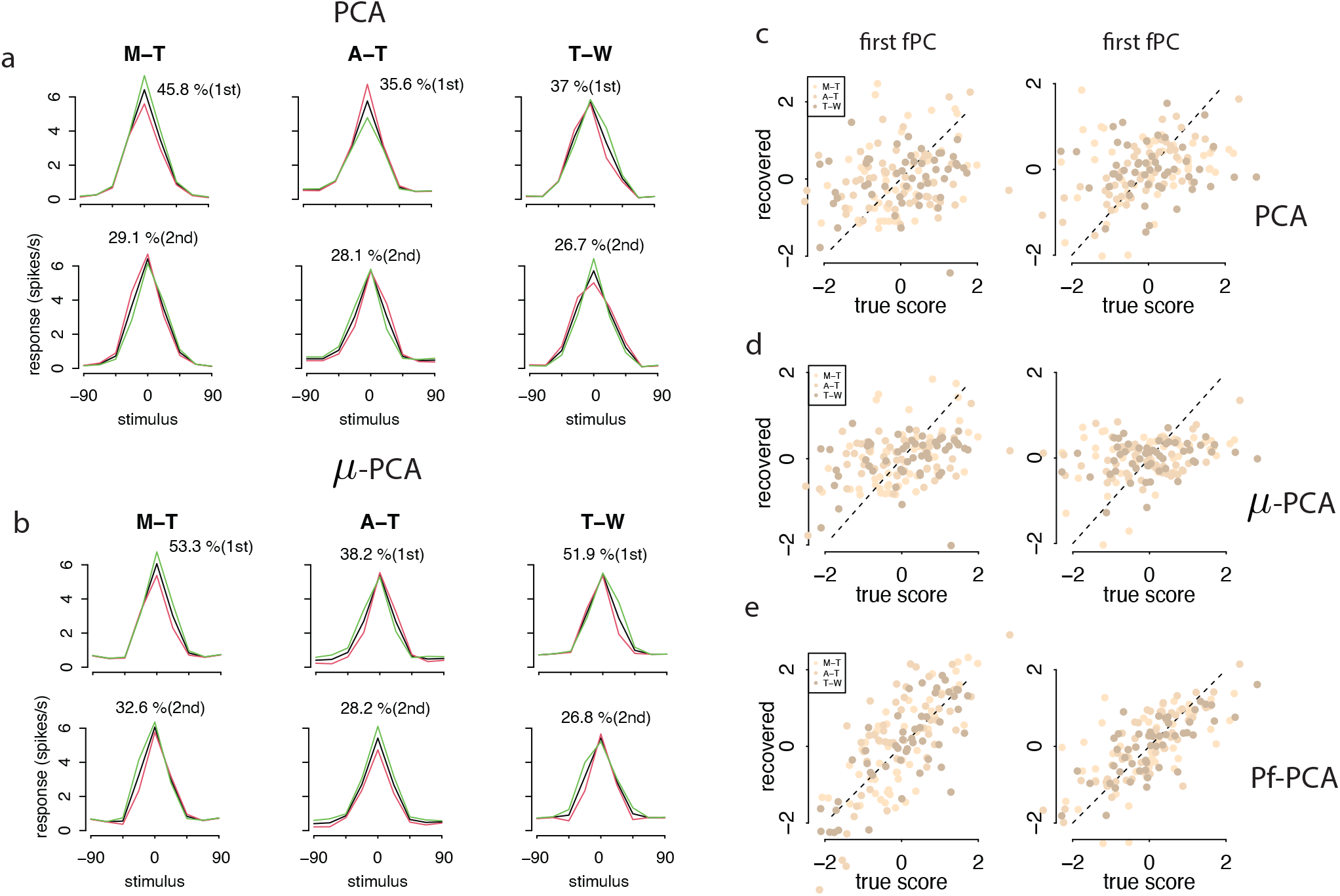
Recovering mixed fluctuations with two components by PCA and *µ*-PCA (related to Fig. S1). (a) Recovering mixed fluctuations with two components by PCA. (b) Recovering mixed fluctuations with two components by *µ*-PCA. (c,d,e). Recovering scores by PCA, *µ*-PCA, and Pf-PCA, respectively. In (a,b), the 1st row denotes the 1st component and the 2nd denotes the 2nd component. In (c,d,e), the 1st scatter denotes the 1st component and the 2nd denotes the 2nd component. In (c), the correlations are 0.22 and 0.31. In (d), the correlations are 0.35 and 0.27. In (e), the correlations are 0.76 and 0.72. M: multiplicative gain; T: tuning shift; W: tuning width change. M-T represents mixing multiplicative gain and tuning shift; same convention for the other two cases. The percentages show the proportions of variance explained by individual component.

### SI.2 Recovering fluctuations for monotonic tuning curves

The response functions of neurons can often be monotonic, e.g., time course in the LIP area [73], response to the motion strength in macaque MT area [98] and time course in the Olfactory bulb [99]. In this section, we address this class of tuning curve shape by using sigmoid functions as a representative example. The fluctuations considered here are the multiplicative gain, the tuning shift, and the slope change (leftmost panel of Fig. S3). Similar to the results reported for bell-shape tuning curves, we found that Pf-PCA could recover the basic structure of fluctuations well (see Fig. S3). It also outperforms the alternative methods (regular PCA and *µ*-PCA).

**Figure S3:**
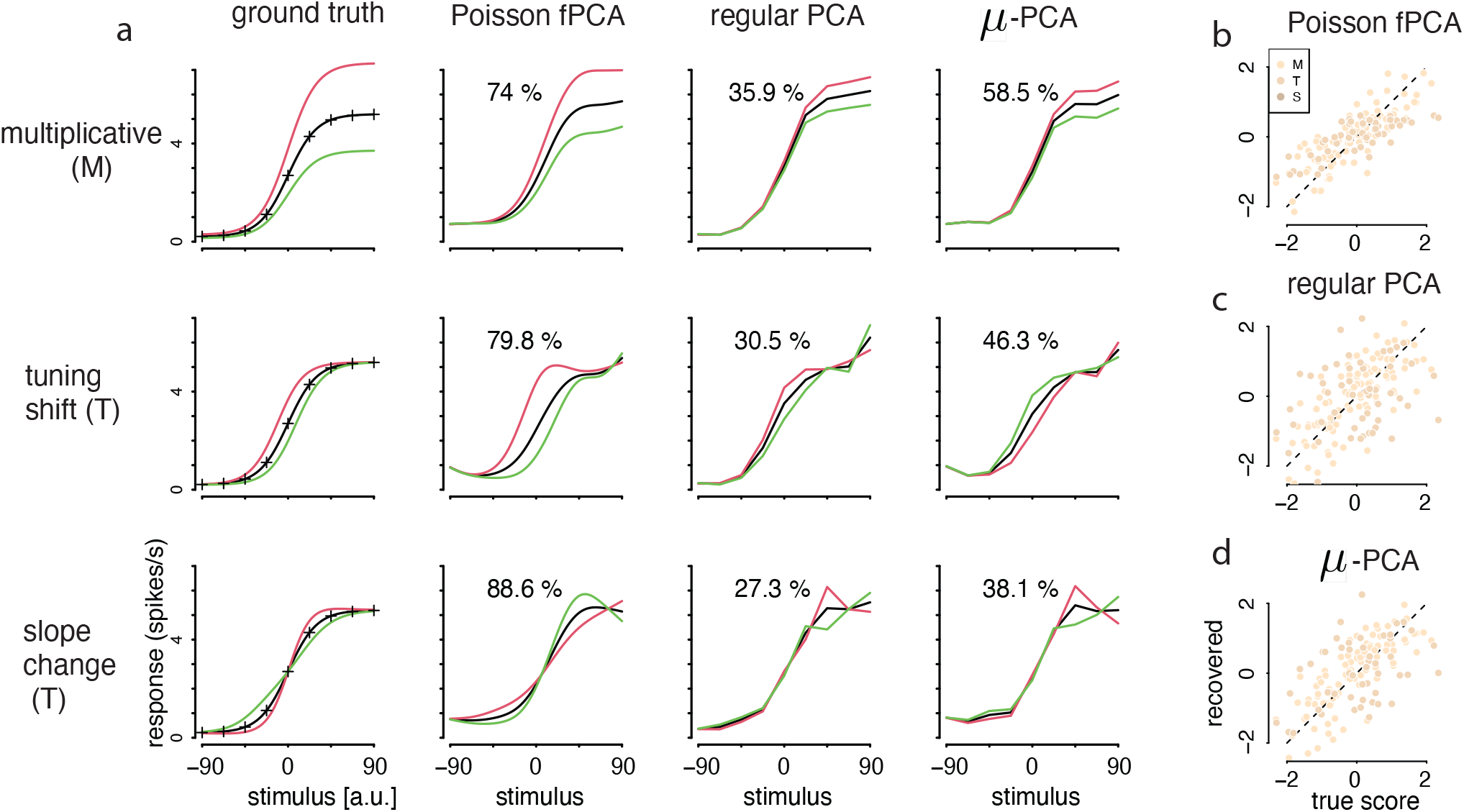
Model validation with monotonic sigmoid response functions (related to Fig. 2 of the Main text). (a) Inferred fluctuations. Same convention as Fig. 2a. Note that in our simulation setting, the first fPC from “perfect” estimation procedure should explain 80% of the variance. (b,c,d) Recovering hidden scores for three methods. Light to dark bisque points denote three fluctuations cases: the multiplicative gain (“M”), the tuning shift (“T”), and the slope change (“S”). In (b,c,d), the correlations are 0.79, 0.43, and 0.45, respectively. Similar to the results for the bell-shape tuning curves, we found that our approach can recover the structure of fluctuations well. Thus our method is not limited by the specific shape of tuning curves.

### SI.3 The first three fPCs are sufficient for accounting most of the variance in the data

We reported the average proportions of variance explained by the first 3 fPCs for each session in Fig.S4.

### SI.4 Recovered slope and intercept from the synthetic dataset

For the purpose of comparing with Fig. 4, we reported the recovered slope and intercept from the synthetic dataset using Pf-PCA in Fig.S5.

**Figure S4:**
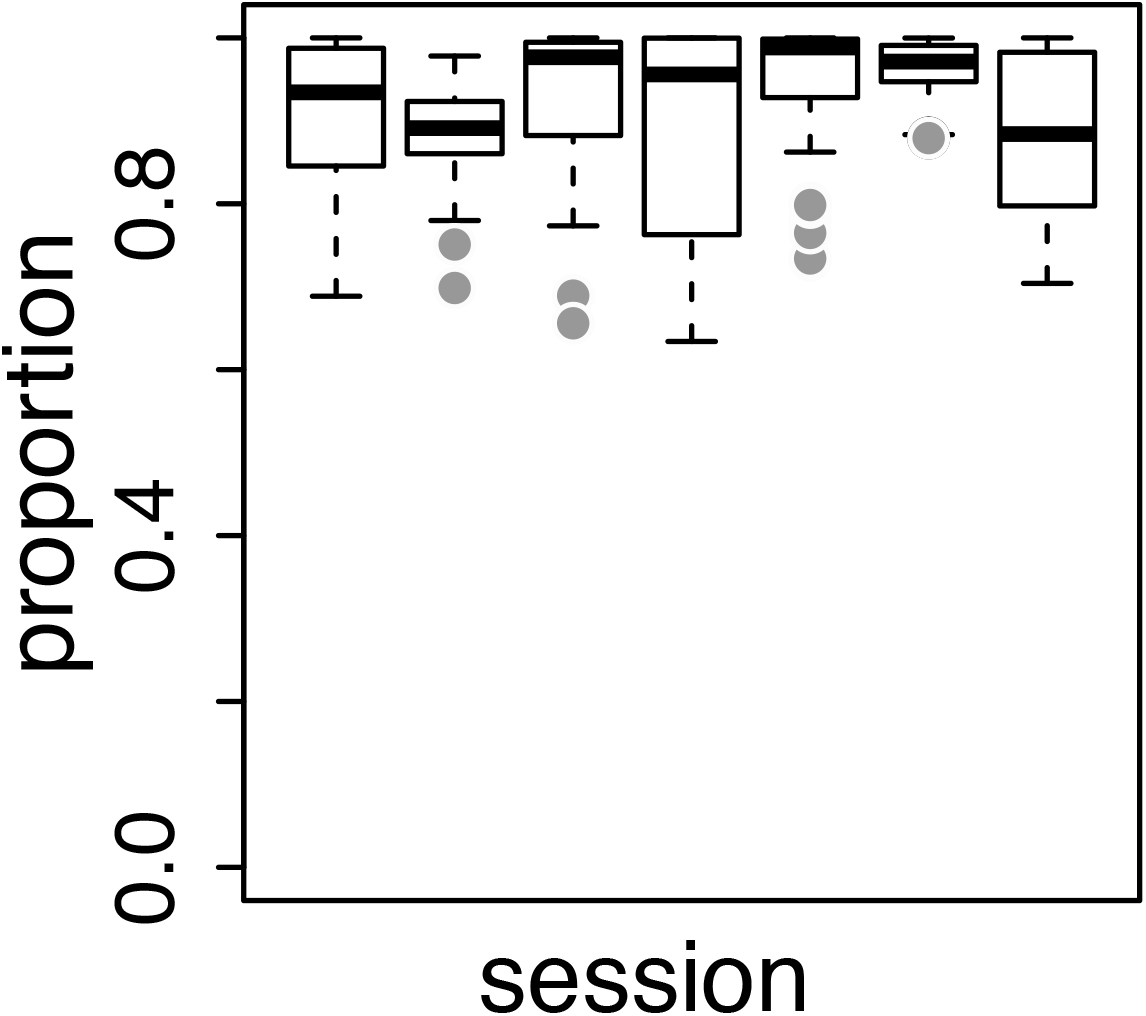
The first three fPCs are sufficient for accounting most of the variance in the data. The average proportion of variance explained from D1 to D7 are 0.91, 0.89, 0.93, 0.90, 0.94, 0.96, and 0.89, respectively.

### SI.5 Population structure of the tuning fluctuations from all 7 sessions

We reported the analysis on the recovered score matrices from all 7 sessions. Fig.S6 listed the results based on real datasets of all 7 sessions. Fig.S7 listed the results inferred from corresponding simulated datasets based on a multiplicative gain model, and Fig.S8 listed the results based on simulated data from an additive gain model.

### SI.6 Further analysis on the FI and 8-way classification task

While our results are consistent with [30] in that both studies found that increased population activities do not lead to substantial increase of population FI, we also noticed some subtle discrepancies between the two studies, because [30] suggested that there was minimal change of population FI when population activity changes. We believe that the difference lie in the difference in the analysis methods. Our results were based on direct estimation of FI, while [30] used 8-way classification by splitting the data into groups with high/low population activity. FI by definition is a local measure of discriminability, thus 8-way classification may not represent an accurate measure of FI. Additionally, by splitting the data into two groups [30], there are likely still substantial amount of tuning fluctuations within each group. Such uncertainty might dominate the classification performance, reducing the amount of change when comparing the high/low groups.

To further understand the potential difference between these two methods, we performed analysis using synthetic data with known ground truth (Fig. 6d). We first generated simulated data which match the summary statistics of the real data, and applied our analysis pipeline to estimate the FI. Next, in a separate analysis, we performed 8-way classifications using the procedure in [30] on the synthetic data. We found that i) our method could reliably estimate the FI, both at the level of individual neurons (Fig. S9) and the neural populations (Fig. 6d); ii) performance based on classification analysis could not be mapped onto the change of FI in obvious ways - higher FI (with decreased population firing rate) does not necessarily leads to better classification performance

**Figure S5:**
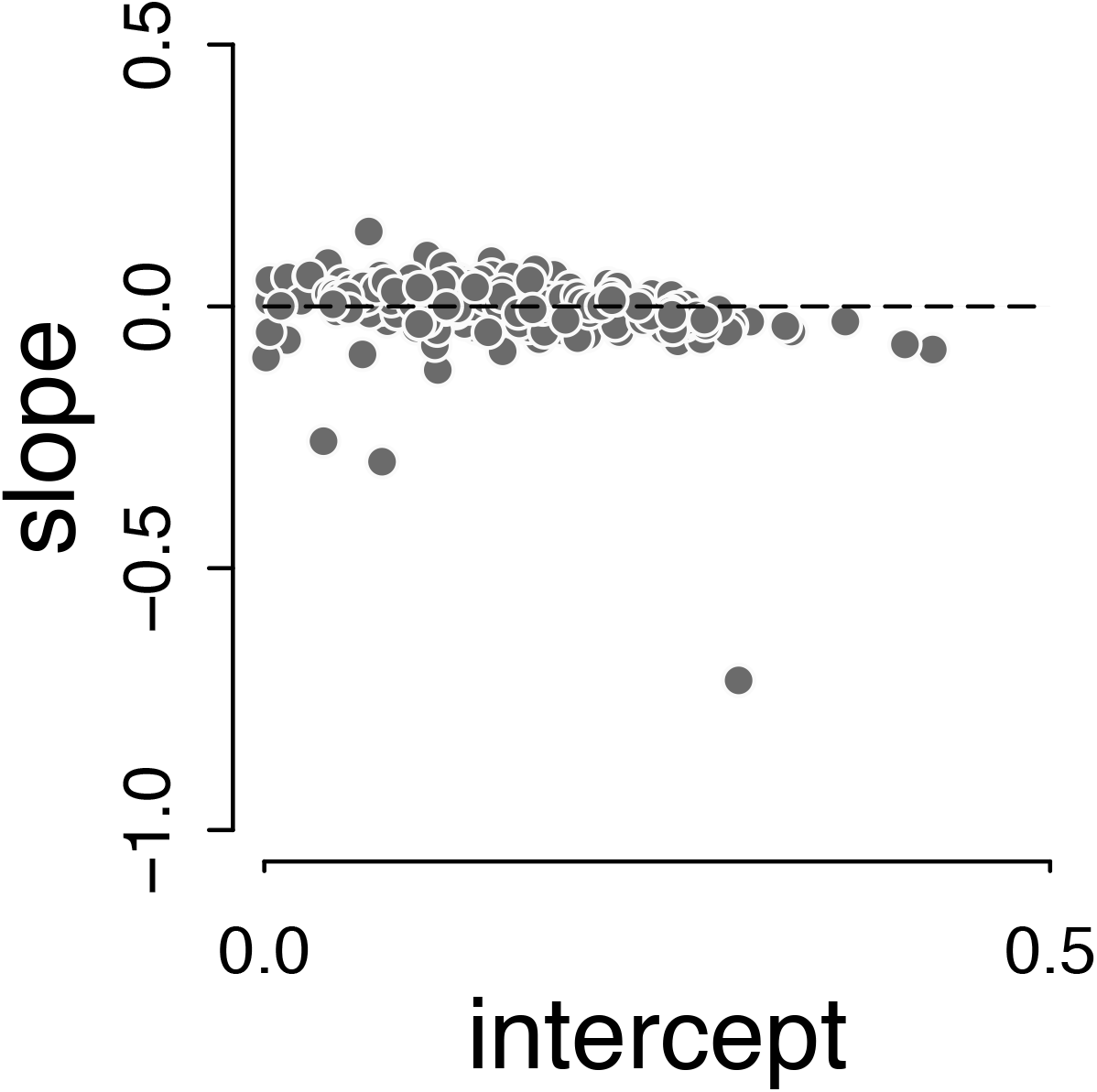
Recovered slope and intercept from the synthetic datasets using Pf-PCA (related to Fig. 4). The synthetic datasets were generated assuming a pure multiplicative gain model while roughly matching individual neuron’s firing rate in the real data. Two observations could be made: first, most of the neurons have a slope that is close to zero as expected; second, the recovered slope is flat over intercept.

**Figure S6:**
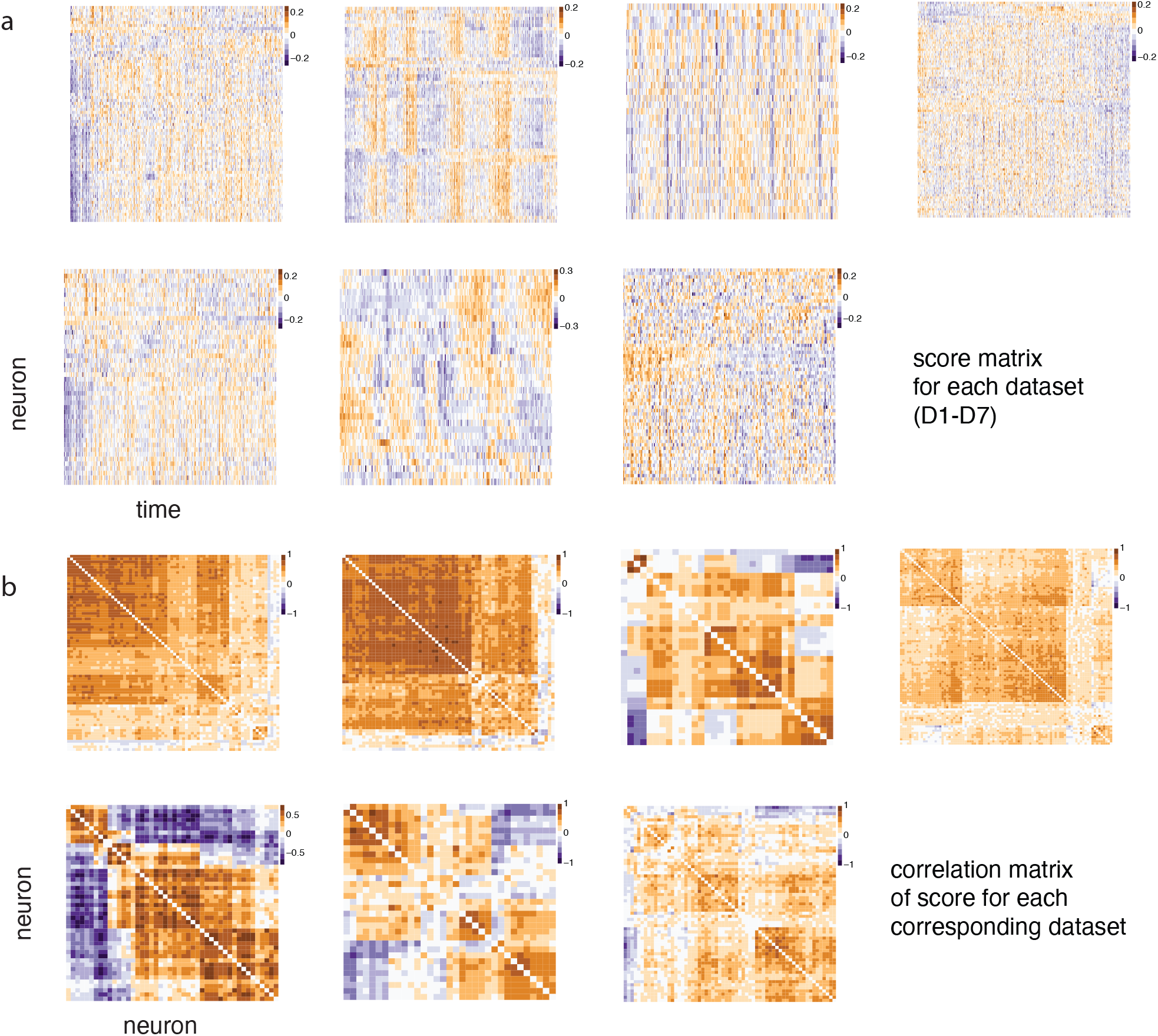
Population structure of the tuning fluctuations in all datasets (related to Fig. 5). (a) The heatmap of scores for individual sessions (from D1 to D7). (b) The corresponding correlation matrix. Neurons were sorted using hierarchical clustering algorithm.

**Figure S7:**
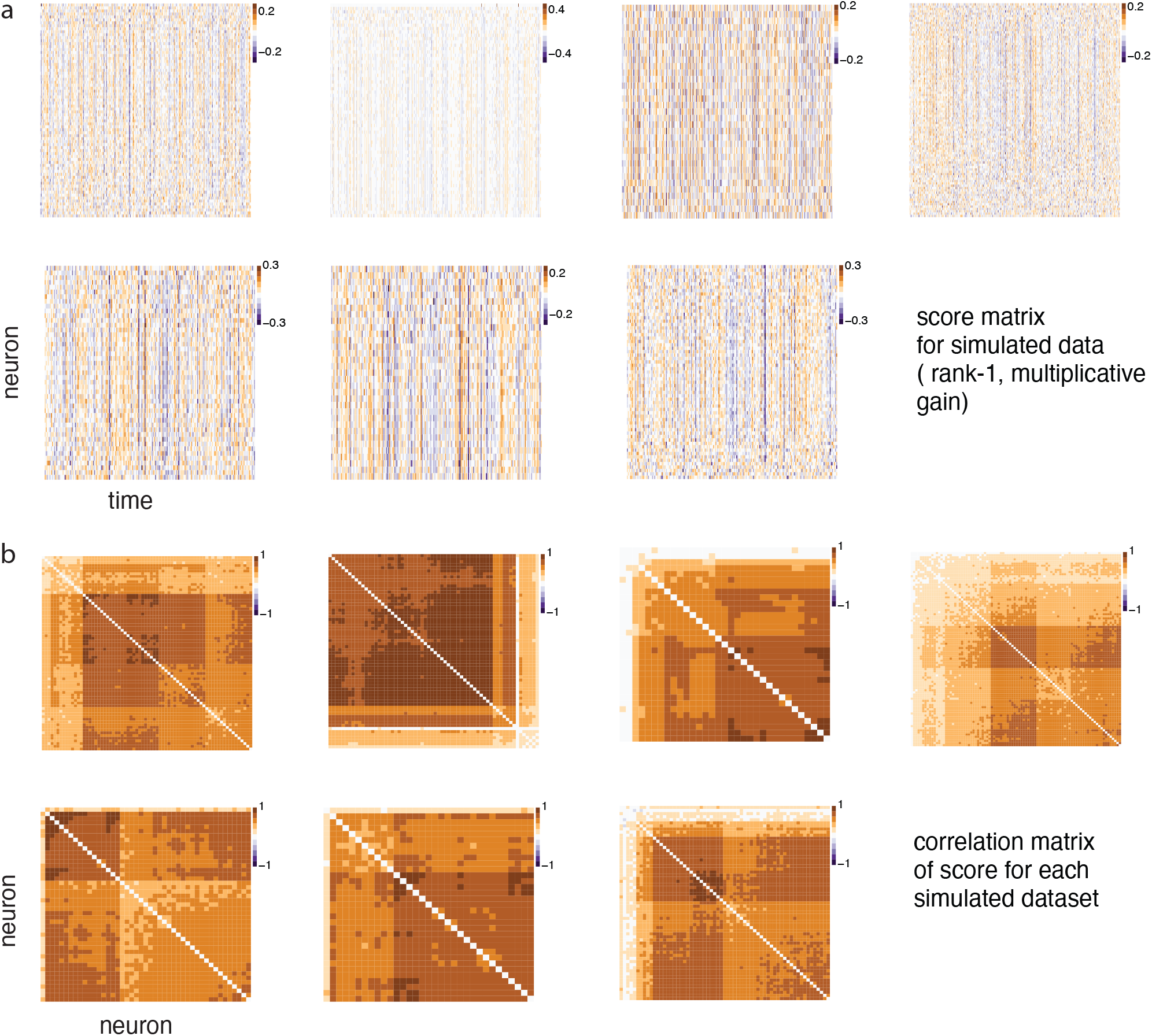
Population structure of the tuning fluctuations in all simulated datasets based on multiplicative gain model (related to Fig. 5). (a)The heatmap of scores inferred from simulated datasets based on multiplicative gain model. (b) The corresponding correlation matrix. Neurons were sorted using hierarchical clustering algorithm.

**Figure S8:**
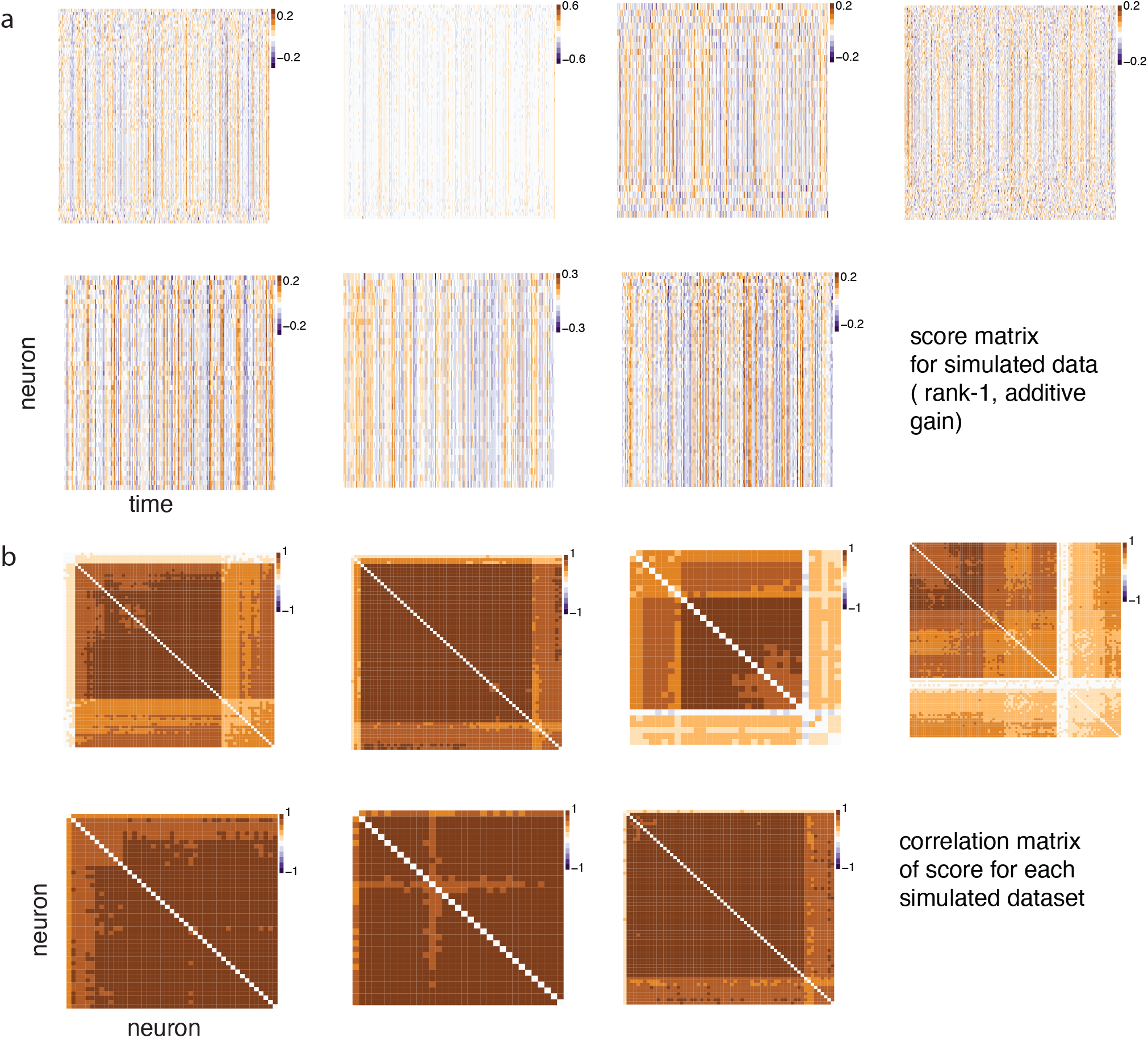
Population structure of the tuning fluctuations in all simulated datasets based on additive gain model (related to Fig. 5). (a) The heatmap of scores inferred from simulated datasets based on additive gain model. (b) The corresponding correlation matrix. Neurons were sorted using hierarchical clustering algorithm.

(Fig. S10). We also verified our classification procedure on the datasets used in [30], and we were indeed able to replicated the classification results in [30] (Fig. S10). These analyses suggest that the two analysis methods may capture different aspects of the code, and generally speaking, 8-way classification task does not lead to accurate characterization of the FI.

### SI.7 Geometry analysis for additive gain model

From comparing with Fig. 7 (a-e), we reported the geometry analysis for an additive gain model in Fig.S11.

**Figure S9:**
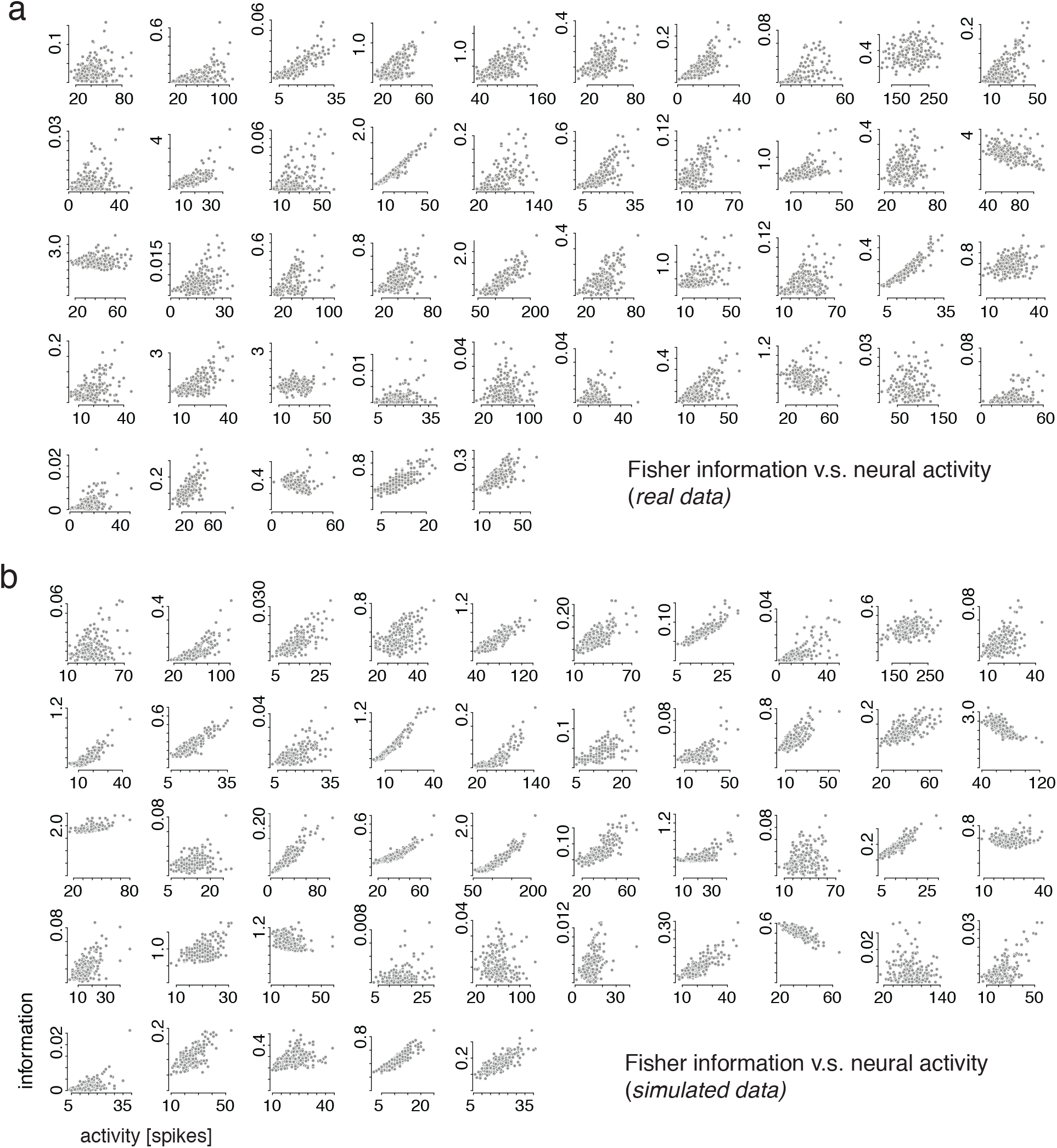
Fisher information as a function of neural activity for individual neurons in one session of data (Session D5). (a) Fisher information as a function of the neural activity for each neuron. (b) Results from the recovery analysis. We simulated count data based on the estimated model by Pf-PCA, then ran our analysis pipeline to estimate the FI. The results suggest that our analysis pipeline is capable of recovering the relationship between the neural activity and the FI in the noise regime similar to the real data.

**Figure S10:**
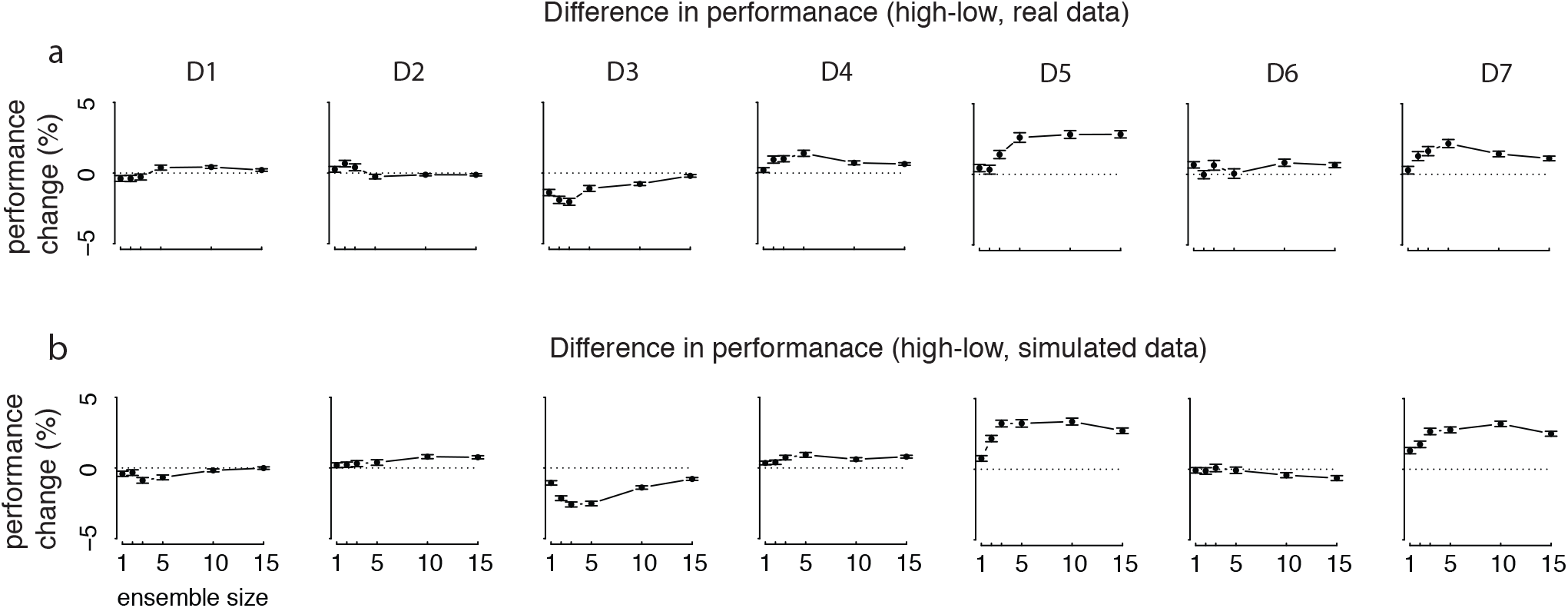
Classification performance as a function of the number of neurons for both real data and simulated data. Error bar: standard deviation of the mean. (a) performance change between high and low trials (high minus low) in each dataset (D1-D7) based on classification procedure described in the Method section. (b) similar to panel (a), but for synthetic data. The results show that classification performance based on the synthetic data recapitulates the pattern in the real data. These observations together with the FI analysis, show that FI and classification performance based on splitting high/low trials as done in [30] do not consistently map onto each other, suggesting the two analysis methods are characterizing non-identical information of the neural code. Note that datasets D1-D4 were from [30]. If one were to combine the performance across these four datasets, the results would show minimal difference between the low and high trials (not shown), consistent with the results reported in [30].

**Figure S11:**
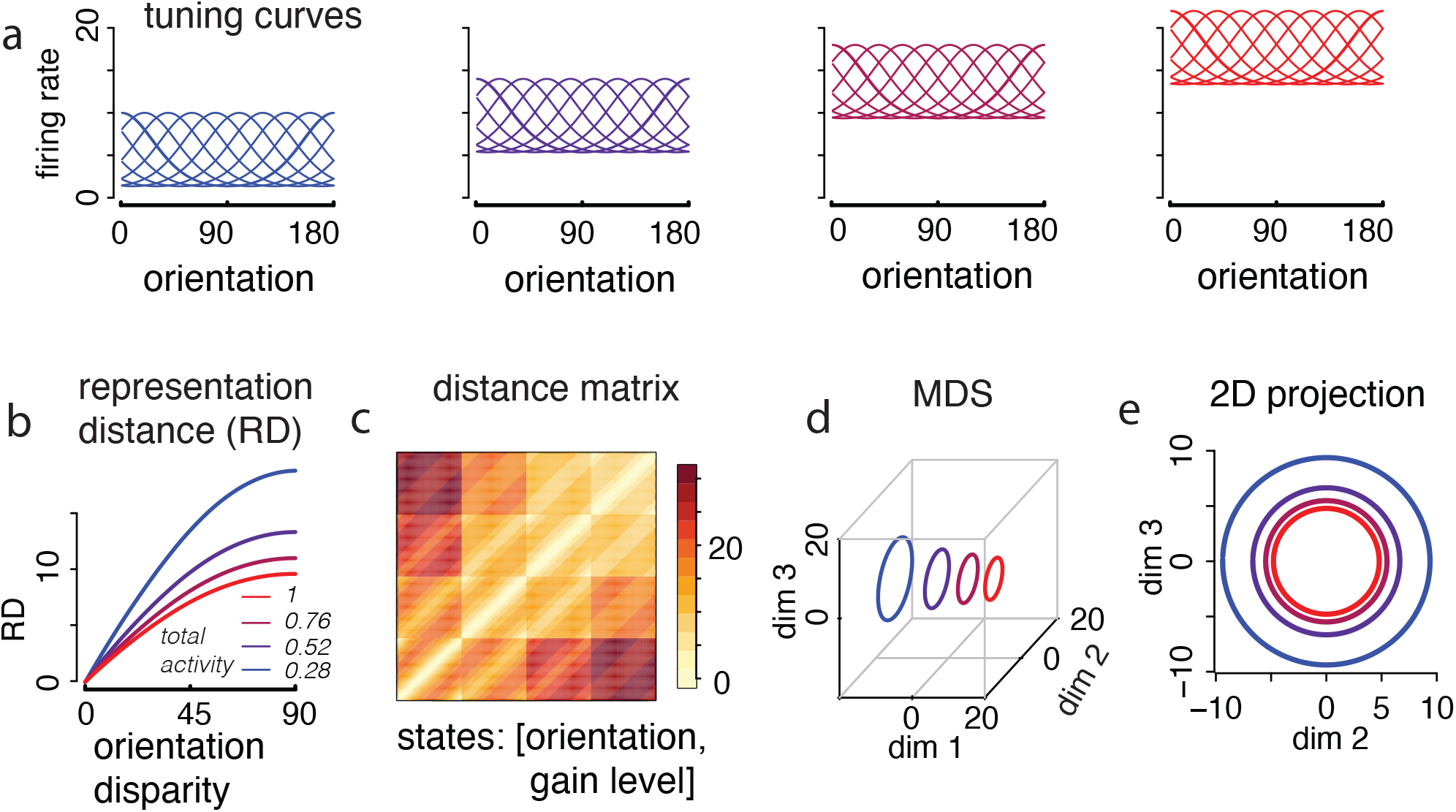
Geometry analysis for additive gain model. (a-e) Similar convention to Fig. 7 (a-e), but for additive gain model. (a) Tuning curves for the population under four different levels of additive gain. (b) The representation distance as a function of the orientation disparity. Inserted: the normalized total activity for each gain level. (c) The representational distance matrix for each pair of states, defined by the orientation and the gain level. We discretized the orientation into 180 bins, results in 180× 4 = 720 states. The states are arranged according to the orientation and states. (d) Results from 3-D MDS. (e) Projection of the 3-D MDS results onto the second and third dimension reveals that the increased activity substantially reduced the size of the representation.

## Notes

### Competing Interest Statement

The authors have declared no competing interest.

